# Ancient loss of catalytic selenocysteine spurred convergent adaptation in a mammalian oxidoreductase

**DOI:** 10.1101/2023.01.03.522577

**Authors:** Jasmin Rees, Gaurab Sarangi, Qing Cheng, Martin Floor, Aida M Andrés, Baldomero Oliva Miguel, Jordi Villà-Freixa, Elias SJ Arnér, Sergi Castellano

## Abstract

Selenocysteine (Sec), the 21^st^ amino acid specified by the genetic code, is a rare selenium-containing residue found in the catalytic site of selenoprotein oxidoreductases. Sec is analogous to the common cysteine (Cys) amino acid but its selenium atom offers physicalchemical properties not provided by the corresponding sulfur atom in Cys. Catalytic sites with Sec in selenoproteins of vertebrates are under strong purifying selection but one enzyme, Glutathione Peroxidase 6 (GPX6), independently exchanged Sec for Cys less than one hundred million years ago in several mammalian lineages. We reconstructed and assayed these ancient enzymes before and after Sec was lost and up to today, and found them to have lost their classic ability to reduce hydroperoxides using glutathione (GSH). This loss of function, however, was accompanied by bursts of amino acid changes in the catalytic domain, with protein sites concertedly changing under positive selection across distant lineages abandoning Sec in GPX6. This demonstrates that when sulfur in Cys impairs catalysis a narrow evolutionary path is followed, with epistasis and pleiotropy leading to convergent evolution and triggering enzymatic properties likely beyond those in classic GPXs. These findings are an unusual example of adaptive convergence towards unexplored oxidoreductase functions during mammalian evolution.

## Main Text

Catalytic residues are largely conserved in enzymes (Sharir-Ivry & Xia, 2021) as they lower the activation energy of reactions and thereby can increase enzymatic turnover. Mutations in these evolutionary constrained active sites typically reduce catalytic activity (Carter & Wells, 1988; Loeb et al., 1989; Rennell et al., 1991), a proxy for fitness in enzymes, and are consequently often deleterious. Still, when such mutations do occur and persist, they allow observation of mutational trajectories that either recover the rate of catalysis (Gromer et al., 2003) or open new protein functions (Jayaraman et al., 2022; Jensen, 1976). These trajectories, however, are limited by the enzyme’s sequence, as epistasis favours mutations whose interactions with other residues compensate for catalytic loss or advance alternative properties, with pleiotropy further limiting trajectories that improve one enzymatic property but compromise another (Storz, 2016; Weinreich et al., 2006). This is best exemplified in orthologous proteins, whose sequence conservation among species provides similar genetic backgrounds to mutations.

Here we study the selenoprotein Glutathione Peroxidase 6 (GPX6_Sec_), focusing on the sporadic replacement in several mammalian lineages of the rare amino acid selenocysteine (Sec) for Cysteine (Cys) (Kryukov et al., 2003), and contrast it to that of proteins of the GPX family using either Sec (GPX1, 2, 3 and 4) or Cys (GPX5, 7 and 8) exclusively. Sec to Cys substitutions across orthologous selenoproteins, as seen in mammalian GPx6_Sec_, are unusual (Castellano et al., 2005), with other GPX_Cys_ proteins emerging only after duplication of a GPX_Sec_ gene (Fig 1a; Mariotti et al., 2012). This is due to the low exchangeability of Sec and Cys in catalysis, likely resulting from the strong purifying selection (Castellano et al., 2009) acting on the lower catalytic activity, lower nucleophilicity and lower efficiency as a leaving group of Cys compared to Sec (Arnér, 2010; Axley et al., 1991; Berry et al., 1992; Johansson et al., 2005; Kim et al., 2015; Lee et al., 2000; Reich & Hondal, 2016)). This suggests that Sec to Cys mutations are deleterious and subject to natural selection (Castellano et al., 2009).

**Figure 1.**
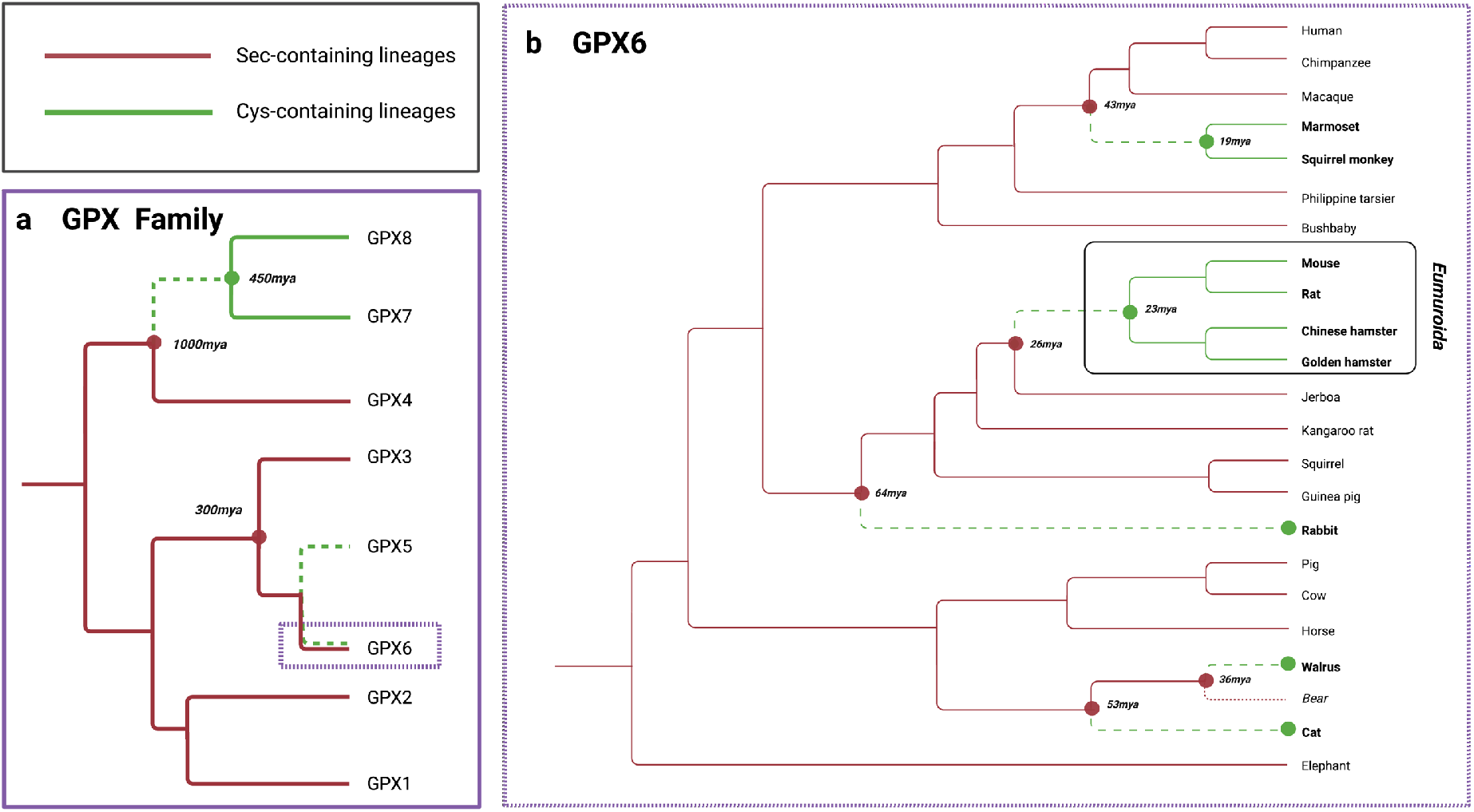
**a.** The phylogeny of the GPX family in Eukaryotes (based on Mariotti et al., 2012), including the dates of the duplications leading to GPX7_Cys_, GPX8_Cys_ and GPX5_Cys_ and their older, single substitutions of Sec to Cys that resulted in enzymes with new properties. **b.** The topology of the phylogeny of the 22 mammals in our analysis. In red, GPx6_Sec_ branches, in green, GPX6_Cys_ ones. Dashed green branches represent GPX6_Cys_ lineages where Sec was lost. Dotted red branch indicates the Bear GPx6_Sec_ lineage, which was not used in the analysis due to sequence quality issues. The GPX6_Cys_ *Eumuroida* clade, a specific group of muroid rodents, is boxed. Approximate ages given by (Chatterjee et al., 2009; Hallström & Janke, 2008; Huchon et al., 2002; Nyakatura & Bininda-Emonds, 2012; Steppan et al., 2004).

GPX_Sec_ activity classically reduces hydroperoxides, particularly hydrogen and lipid peroxides, with glutathione (GSH) as a cofactor. GPX_Cys_ proteins, from early duplications in early mammalian (GPX5_Cys_ from GPX3_Sec_ duplication; around 300 Mya), vertebrate (GPX8_Cys_ from GPX7_Cys_ or GPX4_Sec_ duplication; probably 450 Mya) and metazoan (GPX7_Cys_ from GPX4_Sec_ duplication; more than 1,000 Mya) history (Castellano et al., 2009; Hedges, 2002; Trenz et al., 2021) have evolved a preference for other cofactors, for example thioredoxin in GPX5_Cys_ or protein disulfide isomerase (PDI) in GPX7_Cys_ and GPX8_Cys_ (Nguyen et al., 2011). These Cys-containing proteins act on alternative substrates for peroxidation and may have additional functions, including signalling and oxidative protein folding (Buday & Conrad, 2021; Nguyen et al., 2011; Taylor et al., 2013). Thus, whilst all GPX proteins may protect cells from oxidative damage (Tosatto et al., 2008), those proteins containing Cys may have become imaginative in doing so without being *bona fide* peroxidases, perhaps on account of their lower catalytic turnover. We ask if this is also the case for GPX6_Cys_.

To investigate this, we inferred five independent losses of Sec in GPX6_Sec_ (Fig 1b, dashed green branches) across 22 mammals by reconstructing the ancestral sequence at each node of their phylogeny with PAML (Yang, 2007). These losses occurred in the last 64 million years (Fig 1b, dashed green branches; Chatterjee et al., 2009; Hallström & Janke, 2008; Huchon et al., 2002; Nyakatura & Bininda-Emonds, 2012; Steppan et al., 2004) and have resulted in multiple GPX6_Cys_ lineages. We evaluated the impact of natural selection by calculating independent dN/ ratios (Yang, 2007) for each branch of the mammalian tree (Fig 1 b), including ancestral branches, and found dN/dS ratios in GPX6_Cys_ lineages larger than in neighbouring GPX6_Sec_ lineages, suggesting faster evolution along the branches with Cys (Fig S1). We explicitly tested this hypothesis with a Branch model likelihood ratio test PAML (Yang, 2007)and found that contrasting the dN/dS ratios of GPX6_Cys_ lineages in the branches where Sec was lost (Fig 1b, dashed green branches) to GPX6_Cys_ lineages in the branches inheriting this loss (Fig 1b, solid green branches) and GPX6_Sec_ lineages (Fig 1b, solid red branches), does indeed support a higher dN/dS ratio at the times where Sec was substituted for Cys (LR test; P = 0.002; dN/dS = 0.370 dashed green versus 0.279 solid green versus 0.217 solid red branches in Fig 1b). Because our analysis excluded the Sec to Cys change, a burst of amino acid evolution must have accompanied the loss of Sec.

However, dN/dS inflation across GPX6_Cys_ lineages is still under 1, making it difficult to posit that positive selection is acting during and after Sec loss, rather than relaxed constraint along GPX6_Cys_. The failure of the dN/dS ratio to reach 1 is unsurprising since positive selection acting on GPX6_Cys_ enzymatic properties would mainly impact the catalytic domain, which is otherwise under strong constraint. We thus separately performed the previous likelihood ratio test in its three domains: the N-terminus, the GPX domain and the C-terminus, as defined in the Pfam Database (Mistry et al., 2021). The GPX domain and, to a lesser extent, the C-terminus domain, are essential for the activity of the enzyme, with two (U/C, Q) and two (W,N) key catalytic residues in the GPX and C-terminus domains, respectively, making a catalytic tetrad (Cheng & Arnér, 2017; Toppo et al., 2008; Tosatto et al., 2008), which we found conserved across GPX6_Sec_ and GPX6_Cys_ lineages. In contrast, the N-terminus is not thought essential for catalysis.

As expected, we found the GPX and N-terminus domain of GPX6_Sec_ lineages to be most and least constrained, respectively, based on their dN/dS ratios (Table 1). However, the dN/dS ratio of the GPX domain is, unlike the N- and C-terminus, significantly larger in GPX6_Cys_ lineages at the time Sec was lost (LR test; P = 2×10^-5^; dN/dS = 0.384 dashed green versus 0.186 solid green versus 0.130 solid red branches in Fig 1b). This is again suggestive of evolutionary changes above the overall constraint on the active GPX domain, surrounding the time when Sec is abandoned in catalysis. We then investigated whether this observation was exclusive to GPX6_Cys_, and therefore indicative of rapid evolution associated with the Sec to Cys exchange rather than on the overall antioxidant function of the GPX domain, as others have suggested (Tian et al., 2021). Indeed, when comparing this domain with other enzymes in the GPX family, we found no evidence of dN/dS inflation (Table 1; Table S1) in the lineages where Sec was lost in GPX6 (analogous dashed green branches in Fig 1b in other GPXs), and only a significant inflation in GPX4_Sec_ (Table 1) in the lineages of GPX6 that follow the loss of Sec (analogous solid green branches in Fig 1b in GPX4_Sec_), unrelated to the Sec to Cys exchange. We reason that the dN/dS ratio of the GPX domain in GPX6_Cys_ at the time of Sec loss is unusually large for proteins of the GPX family.

**Table 1.** dN/dS ratios in lineages where GPX6 has Sec (Fig 1b, solid red branches), has gained Cys (Fig 1b dashed green branches) or inherited it (Fig 1b, solid green branches), and the number of identified convergent sites between lineages where GPX6 has gained Cys (Fig 1b, dashed green branches). dN/dS ratios and number of identified convergent sites for the GPX domain in other GPX proteins. The likelihood ratio test contrasts one ratio for all branches (null hypothesis) to different ratios among groups of branches. P-values are obtained from a *χ*^2^ distribution with d.f = 2. *P < 0.05; **P < 0.005; ***P < 0.0005. In bold when significant and accompanied by sites under convergent evolution across GPX6_Cys_ lineages.

| Protein | Region | dN/dS in branches where GPx6 has |  |  |  |  | Convergent sites |
| --- | --- | --- | --- | --- | --- | --- | --- |
|  |  | Sec | Cys when Sec was lost | Inherited Cys | All | P-value | Number |
| GPX6<sub>Cys</sub> | Full length | 0.217 | **0.370** | 0.279 | 0.256 | **0.002\*\*** | **22** |
|  | N-terminus | 0.436 | 0.411 | 0.671 | 0.460 | 0.268 | 3 |
|  | GPX domain | 0.130 | **0.384** | 0.186 | 0.184 | **2x10<sup>-5\*\*\*</sup>** | **12** |
|  | C-terminus | 0.174 | 0.258 | 0.250 | 0.203 | 0.157 | 7 |
| GPX1<sub>Sec</sub> | GPX domain | 0.064 | 0.040 | 0.069 | 0.060 | 0.534 | 0 |
| GPX2<sub>Sec</sub> |  | 0.075 | 0.042 | 0.038 | 0.060 | 0.191 | 0 |
| GPX3<sub>Sec</sub> |  | 0.094 | 0.108 | 0.056 | 0.091 | 0.439 | 1 |
| GPX4<sub>Sec</sub> |  | 0.062 | 0.007 | 0.203 | 0.061 | 1x10<sup>-4***</sup> | 0 |
| GPX5<sub>Sec</sub> |  | 0.233 | 0.145 | 0.219 | 0.212 | 0.227 | 4 |
| GPX7<sub>Cys</sub> | GPX domain | 0.083 | 0.080 | 0.117 | 0.088 | 0.712 | 0 |
| GPX8<sub>Cys</sub> |  | 0.223 | 0.155 | 0.198 | 0.207 | 0.616 | 0 |

We then asked whether this translated into evidence for positive selection on individual sites that accompany the substitution of Sec for Cys in GPX6_Cys_. We contrasted GPX6_Cys_ lineages for this domain at the time of Sec loss (Fig 1 b, dashed green branches) with all other lineages, and found a branch-site model likelihood ratio test (PAML; Yang, 2007) significant (LR test; P = 0.046; Table S2). This is suggestive of sites under positive selection in the GPX domain when Sec was lost. Since epistasis and pleiotropy can limit the mutations that are favoured in a particular genetic background, we also asked if convergence had occurred in the lineages where Cys was lost. We identified convergent sites as those having changed to the same amino acid or simply repeatedly changed with CONVERG2 (Zhang & Kumar, 1997), and found that convergence between lineages where Sec was lost (Fig 1b, dashed green branches), was highest (Table S3).

Moreover, most convergent sites are found in the GPX domain, and the least in the N-terminus (54.6% in the GPX domain, followed by 31.8% and 13.6% in the C-terminus and N-terminus respectively). Convergence is largely subdued in the GPX6_Cys_ lineages inheriting the loss of Sec (Fig 1b, solid green branches) and minimal in the GPX6_Sec_ lineages and other GPX_Sec_ and GPX_Cys_ enzymes (Tables S3-10; Fig S2). Further, simulations of protein evolution incorporating the accelerated rate of amino acid change in GPX6_Cys_ sequences, including the GPX domain, cannot reproduce the pattern of convergence observed between these lineages at the time of Sec loss (Seq-gen; Fig S5-6). However, these simulations show that the few weak convergence signatures in other GPXs are as expected (Fig S7-13), and we presume they result from chance. We conclude that signatures of convergence are exclusive to GPX6_Cys_ and strongest in its GPX domain at the time that Sec was lost, acting under selective pressure and possibly epistasis.

We observed that the highest level of convergence is between the basal *Eumuroida* (Fig 1b, dashed green line in box) and its genetically closer GPX6_Cys_ lineages, particularly the rabbit (Fig S2; Table S3). We therefore examined the 25 sites at the root of the *Eumuroida* (Fig 2a, dashed green branch) that changed along Sec and found that 14 of them bear signatures of convergence across the GPX6_Cys_ lineages (Fig 2a, green box). These convergent sites are, again, mainly in the GPX catalytic domain, 64.3% of them (Table S2), and are enriched for the signatures of positive selection that we observe along branches leading to *Eumuroida*, rabbit and marmoset-squirrel monkey (M-W U test, P = 1.573e-7). We also find signatures of positive selection, albeit weaker, in convergent sites in the GPX6_Cys_ lineages following the loss of Sec (M-W U test, P=0.007) (Fig 2a, solid green branches) but not preceding it, in agreement with adaptive convergence acting on the Sec to Cys exchange.

**Figure 2.**
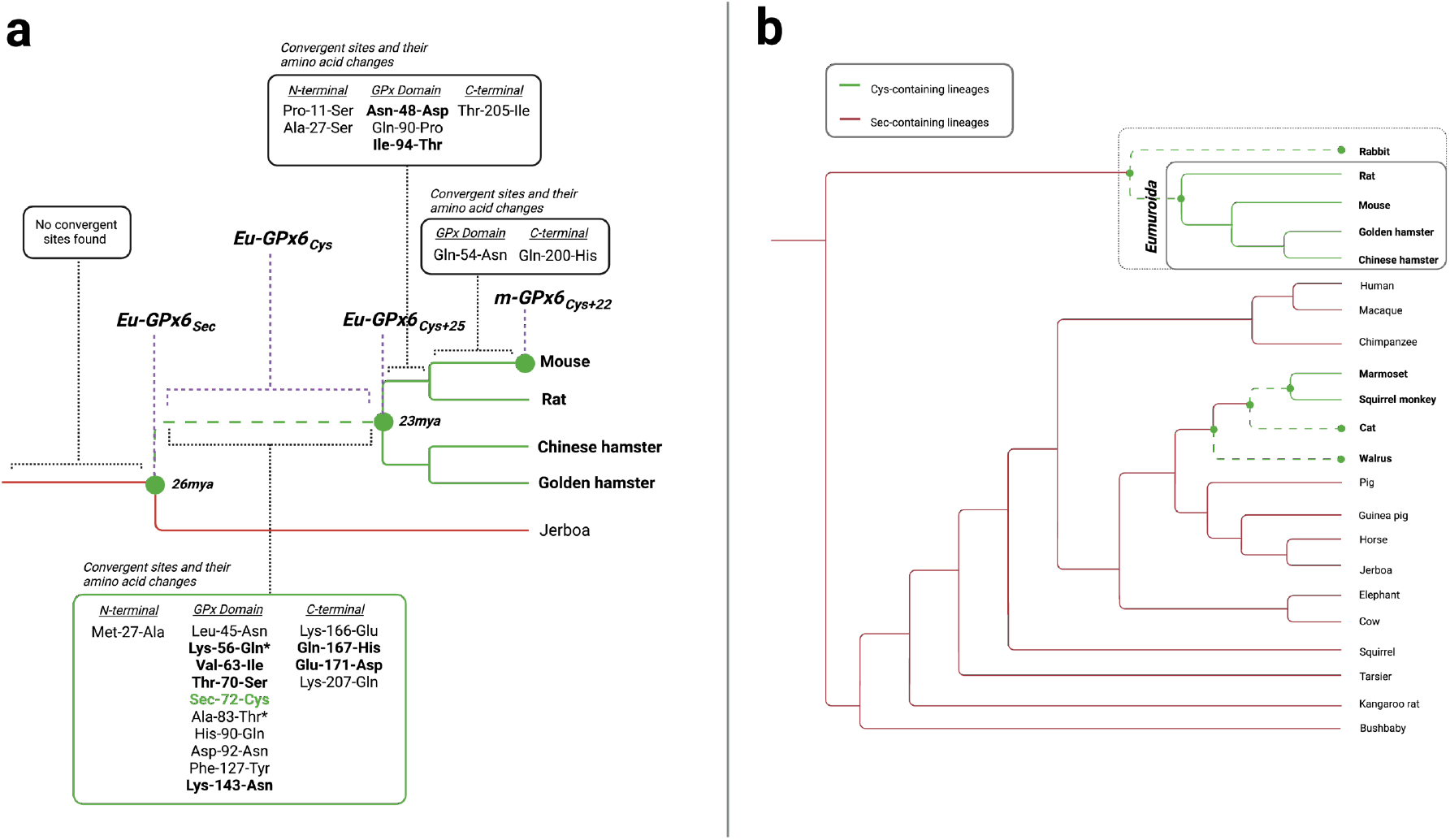
**a**. Topology of the phylogeny of the *Eumuroida* GPX6_Cys_ clade, green branches, with the Jerboa GPX6_Sec_ lineage, red branch, as an outgroup. The basal *Eumuroida* lineage, dashed green branch, abandoned Sec in catalysis for Cys with a burst of 25 additional amino acid changes 23-26 million years ago. 14 of these 25 changing amino acid sites, plus the Sec to Cys site, have signatures of convergence (CONVERG2; Zhang & Kumar, 1997) across GPX6_Cys_ lineages (Fig 1b, green branches). These sites are located in the N-terminal (1 site), GPX domain (10 sites, including Sec to Cys) and C-terminal (4 sites) domains and have repeatedly changed in the GPX6_Cys_ lineages towards similar or the same (in bold) amino acid (green box). No convergent sites are found immediately before the loss of Sec and only a handful immediately after. Further, the * denotes sites with a posterior probability of positive selection in the upper 90^th^ percentile across the GPX domain in GPX6_Cys_ lineages, which are significantly enriched at the time Sec was abandoned. **b**. Topology of the phylogenetic tree, with midpoint rooting, from the 14 convergent sites accompanying the Sec to Cys substitution (Fig 2a, green box) in the basal *Eumuroida* GPX6_Cys_ lineage (Fig 2a, dashed green branch). In sharp contrast to the species phylogeny (Fig 1b), the GPX6_Sec_ lineages now form two clades.

Because adaptive convergence can mimic shared ancestry, it may distort the topology of the species phylogeny (Fig 1b) and, indeed, we found that a tree reconstructed (PhyML; Guindon et al., 2010) from the GPX domain pulls the rabbit lineage closer to the *Eumuroida* clade (Fig S3d). This, to a lesser extent, is also observed with the C-terminus (Fig S3e), which contributes to catalysis, but is not observed with the N-terminus domain (Fig S3c) nor with other GPX proteins (Fig S4). The strongest departure from the species tree is reconstructed from the 14 convergent sites changing at the root of the *Eumuroida* 23-26 million years ago (Huchon et al., 2002), plus the Sec to Cys substitution (Fig 2b). In this tree, despite their large divergence, the GPX6_Cys_ species form two clades. One clade shows the rabbit lineage sharing a most recent common ancestor, to the exclusion of all other species, with the 64 million years apart *Eumuroida*, and the other clade groups the remaining 100 million years apart GPX6_Cys_ lineages (Hallström & Janke, 2008). This supports a role for convergence (Edwards, 2009) in driving adaptive changes, perhaps compensating for loss of enzymatic activity, or opening new properties, as it seems to have happened with GPX5_Cys_, GPX7_Cys_ and GPX8_Cys_ enzymes that lost Sec much earlier (Fig 1a) (Chen et al., 2016; Herbette et al., 2007).

Given these signatures of adaptive convergence in the *Eumuroida*, we explored the functional consequences of such changes. We reconstructed three ancient proteins existing 23-26 million years ago at the root of this clade (Fig 2a, dashed green branch) and assessed them experimentally and computationally. We also included a fourth modern protein within this analysis. These proteins are: 1) the ancestral protein before the loss of Sec, Eu-GPX6_Sec_, taken from the common ancestor of the *Eumuroida* and Jerboa species 26 million years ago (Huchon et al., 2002); 2) the same ancestral protein with Cys instead of Sec, Eu-GPX6_Cys_; 3) the ancestral but later-day protein with Cys and 25 other amino acids changes, Eu-GPX6_Cys+25_, taken from the common ancestor of the *Eumuroida* species 23 million years ago (15 of 26 these amino acid changing sites, including the Cys site, have signatures of adaptive convergence; Fig 2a); and 4) the present-day mouse protein, m-GPX6_Cys+22_, with 22 additional amino acid changes (19 substitutions and a 3 C-terminal extension) from Eu-GPX6_Cys+25_ and no clear signatures of adaptive convergence (Fig 2a). In the latter protein, we also mutated the enzyme to contain either Sec or redox inactive serine (Ser) for comparisons of activity with the Sec- and Cys-variants.

We reconstructed these ancient and modern proteins and produced them as recombinant proteins heterologously expressed in *Escherichia coli*. The Sec insertion system in bacteria is non-compatible with mammalian selenoprotein-encoding genes, hampering the production of proteins with Sec; we thus employed a method we recently developed utilizing UAG redefined as a Sec codon in a release factor-1 deficient *E. coli* host strain lacking other UAG codons (Cheng & Arnér, 2017). We first compared the catalytic activity of Eu-GPX6_Sec_ and Eu-GPX6_Cys_ with H_2_O_2_ as the peroxide substrate and GSH as the reducing agent, with the expectation that substitution of Sec for Cys would lower its turnover (Axley et al., 1991; Berry et al., 1992; Johansson et al., 2005; Kim et al., 2015). Indeed, the ancient Eu-GPX6_Sec_ protein displays the classic peroxidase activity of Sec-containing GPX enzymes, whereas Eu-GPX6_Cys_, had almost no activity for this reaction (Fig 3a).

**Figure 3.**
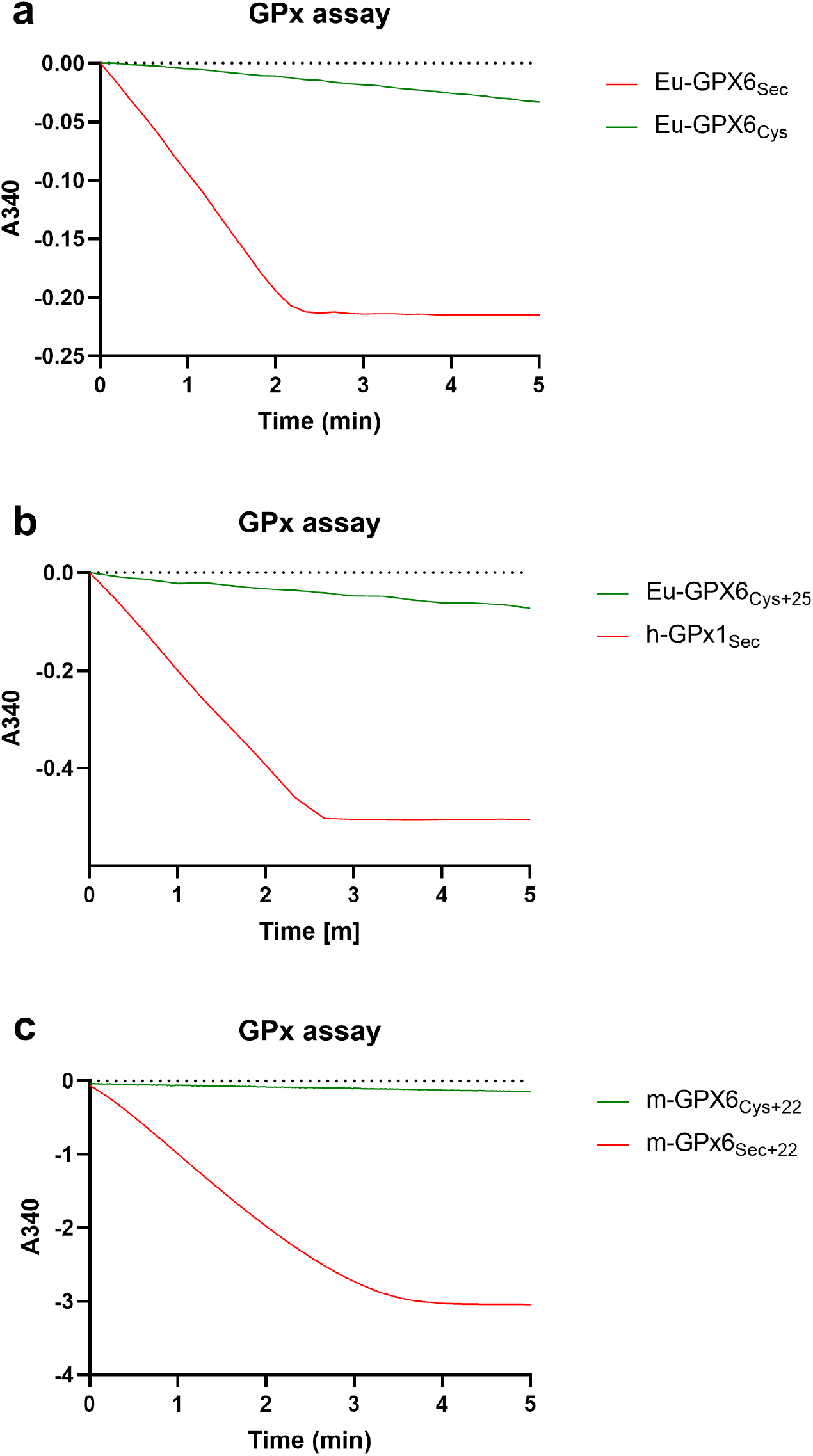
**a**. Experimental assessment of peroxidase reaction with H_2_O_2_ as a substrate for ancient Eu-GPX6_Sec_ (red) and Eu-GPX6_Cys_ (green). NADPH consumption by GR is indicated by the decrease in absorbance at 340 nm over time in the coupled assay (see Methods section for further details). **b**. Equivalent assay for ancient Eu-GPX6_Cys+25_ (green), which has very limited activity compared to human GPx1 (red) used here as a positive control. **c**. Equivalent assay for modern m-GPX6_Cys+22_ (green), again with scant activity, which is recovered once this protein is mutated to contain Sec, m-GPX6_Sec+22_ (red).

The large drop in catalysis from Eu-GPX6_Sec_ to Eu-GPX6_Cys_ coincides with signatures of convergent adaptive evolution along the basal *Eumuroida* lineage (Fig 2a), questioning the role of the accompanying amino acid changes to the loss of Sec. We thus aimed to test the 25 additional changes along the basal *Eumuroida* lineage by measuring catalytic activity in Eu-GPX6_Cys+25_ on H_2_O_2_ with GSH. Remarkably, with this enzyme variant classic GPX activity was not recovered (Fig 3b).

We then turned to the extant m-GPX6_Cys+22_ protein (Fig. 2a), which is 90% identical to Eu-GPX6_Cys+25_, and expressed in the mouse embryo, testis, olfactory epithelium and brain (Goltyaev et al., 2020; Kryukov et al., 2003; Shema et al., 2015). When knocked down, this modern mouse protein has a neurological phenotype (Shema et al., 2015). Interestingly, this Cys-containing variant also lacks classic GPX activity with H_2_O_2_ and GSH (Fig 3c), again suggesting that adaptive changes in the evolution of this protein do not simply act to recapitulate Sec activity. However, classic GPX activity was re-acquired when Cys was mutated back into Sec, producing the synthetic m-GPX6_Sec+22_ variant (Fig 3c). Indeed, our computational analysis suggest that the binding of GSH and overall structures of the enzymes (Fig 4a) have not been adversely affected by the acquisition of Cys and that convergent amino acid substitutions are mainly located in the enzyme’s surface (Fig 4b). This is also the case for GPX6 in other lineages losing Sec (Fig S14).

**Figure 4.**
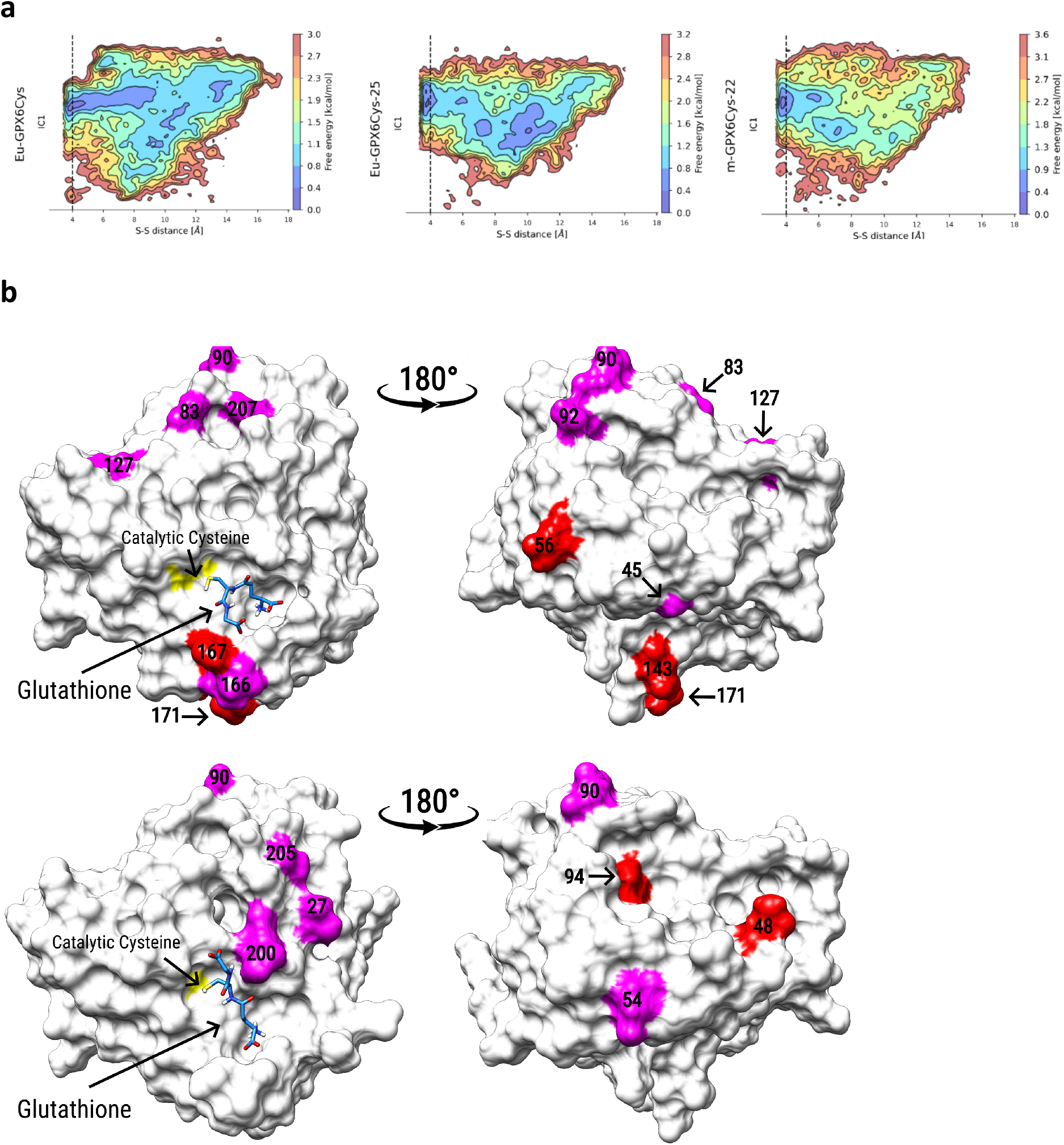
**a**. Free energy profiles for the docking of glutathione to Eu-GPX6_Cys_ (left), Eu-GPX6_Cys_+25 (centre) and m-GPX6_Cys_+22 (right). The x-axis represents the distance between the catalytic cysteine sulphur atom and the ligand’s sulphur atom, while the Y-axis shows the slowest TICA coordinate (IC1). The vertical dashed line represents a 4Å distance, with the free energy minimum in the three enzymes within this reactive catalytic distance. **b**. Convergence patterns (Fig 2a) from Eu-GPX6_Cys_ to Eu-GPX6_Cys_+25 (top) and from Eu-GPX6_Cys_+25 to m-GPX6_Cys_+22 (Mouse-GPX6) (bottom). Sites converging towards similar (magenta) or the same (red) amino acids are shown with their sequence position. The catalytic cysteine (yellow) is shown with the glutathione best binding energy conformation (green) sampled during docking simulations.

It follows that substituting Sec for Cys in GPX6, and thereby abandoning selenium for sulfur, leads to amino acid changes following a narrow evolutionary path, one limited by convergence and the preservation of enzymatic properties expected to be important for overall antioxidant activity. This was likely driven by epistasis from conserved catalytic sites (Sharir-Ivry & Xia, 2021), and perhaps pleiotropy. Furthermore, while GPX6_Cys_ proteins across mammals (Fig S14) appear to be able to recover their classic GPX function with Sec, there exists the possibility that its loss has resulted in subtly different properties in an enzyme now apparently devoid of its classic function. Only comprehensive functional characterizations of these individual GPX6_Cys_ enzymes in mammals will provide insights into their current roles. Thus, similar to old and single losses of Sec occurring hundreds of millions of years ago in GPX5_Cys_, GPX7_Cys_ and GPX8_Cys_ enzymes (Trenz et al., 2021), we suggest that GPX6_Cys_ proteins have also gained yet unidentified abilities, though acquired more recently and independently across lineages.

In conclusion, we present the first evidence for molecular convergence of changes in proteins when abandoning unusual selenium in catalysis for common sulfur. These concerted changes follow a certain path, maintaining some enzymatic properties and possibly adding new ones. Because multiple non-vertebrate species have completely abandoned enzymatic selenium for sulfur, we wonder whether other convergent adaptations leading to uncharted functions remain hidden in nature.

## Methods

### GPX6 and other GPX sequences

The GPX6 coding sequences and proteins for 22 present-day mammal species were obtained from SelenoDB 2.0 (Romagné et al., 2014)now available at <u>selenodb.crg.eu</u>, and Ensembl (Yates et al., 2020), chosen from their availability and breadth across the mammalian tree. These mammals were: Chimpanzee (*Pan troglodytes*), Human (*Homo sapiens*), Macaque (*Macaca mulatta*), Marmoset (*Callithrix jacchus*), Bolivian squirrel monkey (*Saimiri boliviensis*), Tarsier (*Carlito syrichta*), Bushbaby (*Otolemur garnettii*), House mouse (*Mus musculus*), Rat (*Rattus norvegicus*), Chinese hamster (*Cricetulus griseus*), Golden hamster (*Mesocricetus auratus*), Lesser Egyptian jerboa (*Jaculus jaculus*), Kangaroo rat (*Dipodomys ordii*), Squirrel (*Ictidomys tridecemlineatus*), Guinea pig (*Cavia porcellus*), Rabbit (*Oyctolagus cuniculus*), Walrus (*Odobenus rosmarus*), Cat (*Felis catus*), Horse (*Equus caballus*), Cow (*Bos taurus*), Pig (*Sus scrofa*) and Elephant (*Loxodonta africana*).

The Ensembl species tree (available at www.ensembl.org) was used as the topology of the phylogeny of these mammals, with the exception of the walrus which was added according to various additional sources (Higdon et al., 2007). These species include nine mammals where GPX6 contains Cys instead of Sec (Fig 1).

The orthologous GPX6 coding sequences and proteins were aligned using MAFFT (Katoh et al., 2019). The posterior probability of each individual aligned position was then calculated using a modified version of HMMER (Potter et al., 2018). In short, each protein multiple alignment was first converted into a Hidden Markov Model before the use of a forwardbackward algorithm (Durbin et al., 1998) to perform posterior decoding. The calculated posterior probability integrates the uncertainty of the alignment around an aligned position, representing our degree of confidence in each individual aligned protein residue or gap in a multiple alignment.

Positions with an average posterior probability below 0.95 were removed from input to CODEML from PAML (Yang, 2007)owing to concerns of misalignment. The untrustworthy positions are generally found where gaps create alignment uncertainty or where sequence divergence contributes to alignment uncertainty. However, our probabilistic approach allowed us to keep those regions of the alignment with gaps and amino acid differences that are nevertheless confidently aligned in the multiple alignments.

The coding sequences for other members of the GPX family, where all species have Sec (GPX1,2, 3 and 4) or Cys (GPX5, 7 and 8), were also obtained from SelenoDB 2.0 (Romagné et al., 2014) or, if not available, from Ensembl(Yates et al., 2020), and aligned as described above.

### Inferring the Loss of Sec in GPX6

The ancestral sequences of GPX6 for our set of 22 mammals were reconstructed using their present-day sequences and our phylogenetic tree of GPX6 (Figure 1) using the PAML package (Yang, 2007). This package inferred the sequence of all ancestral nodes across the mammalian tree and was used to infer the independent losses of Sec throughout the mammalian lineage. The independent losses of Sec within the walrus and cat lineages were inferred according to the most parsimonious scenario when accounting for the presence of Sec in the Ursidae lineage (included as bear in Fig 1). The approximate ages of lineages with Sec loss are given as ranges in Fig 1, as collected from various sources describing split times in the mammalian phylogeny (Chatterjee et al., 2009; Hallström & Janke, 2008; Higdon et al., 2007; Huchon et al., 2002; Nyakatura & Bininda-Emonds, 2012; Steppan et al., 2004).

Further to the PAML inference, the ancestral sequences were also inferred using two additional programs: Ancestor v1.1 (Diallo et al., 2010) and FastML (Moshe & Pupko, 2019). FastML had the options of using either amino acid or nucleotide sequences of present-day species as input to infer the sequences, whereas Ancestor v1.1 (and PAML) only uses the nucleotide sequences for inference. Using both FastML input methods, this gave us four inferred sequences for all ancestral nodes. The four inferred sequences were then aligned using MAFFT (Katoh et al., 2019) and the residue with most support was taken as the consensus residue for each site.

### dN/dS ratios in GPX proteins

The dN/dS ratio was computed using the CODEML package from PAML (Yang, 2007), using the aligned GPX6 coding sequences and established tree topology (Fig 1) as input. This dN/dS is used as a quantification of the strength of selection acting on proteins, where dN is the rate of non-synonymous substitutions per non-synonymous sites and dS is the rate of synonymous substitutions per synonymous sites. Finally, the UGA codon encoding the Sec amino acid was considered an ambiguity character and, hence, not included in the dN/dS calculation. This makes our tests conservative when comparing the patterns of evolution in proteins that have lost Sec and gained Cys to those that have not.

Independent dN/dS ratios for each branch were estimated using the free-ratio model (model = 1) in PAML, which allows the dN/dS ratio to vary amongst the branches of the phylogenetic tree. This was used to compare the rate of evolution in the lineages that retain Sec and those that have exchanged Sec for Cys. Given this preliminary comparison, the CODEML branch model (model = 2) was then used to explicitly test our hypothesis of a faster rate of evolution in lineages where Sec was lost.

The CODEML branch model (model = 2) which allows us to specify the number of independent dN/dS ratios and on which branches they lie. We used this model to compare the dN/dS ratios between three groups of branches: the branches with Sec (Fig 1; solid red branches), the branches where Sec is exchanged for Cys (Fig 1; dashed green branches) and the branches where Cys is maintained (Fig 1; solid green branches). By using this comparison, we compare if the dN/dS ratio was significantly different in lineages at the time surrounding the loss of Sec compared to lineages where Sec was not lost, or where Cys was maintained and presumably any fitness reduced as a result of the loss of Sec had been recovered.

The branch models were compared to the null model (M0 model, model = 0) where singular dN/dS values are estimated for all branches. The likelihood of each of the two models were then compared to give a likelihood ratio, comparing if three dN/dS ratios across the tree is a better fit than the null model of a singular dN/dS ratio across all branches. The likelihood ratio was used to calculate the significance of the difference in fit between the two models in the form of a p-value (shown in Table 1). A significant deviation between the two models indicates a difference between the dN/dS ratios across the three groups of branches in the tree.

This analysis was repeated for other genes in the GPX family, comparing the rates of evolution of the lineages that have lost Sec in GPX6 and those that have not lost Sec or have maintained Cys (again with CODEML, branch model = 2) with the sequences of the Sec-containing GPX proteins (GPX1, 2, 3 and 4) and the Cys-containing proteins (GPX7, and 8). The likelihood ratio of the branch model to the null model was again calculated to give a P-value for significance (Table 1).

### dN/dS Ratios in the Protein Domains of GPX6

To further explore the rate of evolution over the GPX6 protein, we separated the protein into three domains: N-terminus, GPX domain and C-terminus, as defined in the PFAM (Mistry et al., 2021). Here, the GPX domain is essential for the catalytic activity of the enzyme with the C-terminus believed to also contribute to catalytic function (Toppo et al., 2008). The dN/dS ratios for each domain were then compared between the Sec-containing lineages, the lineages where Sec was exchanged for Cys and the lineages where Cys was maintained of the GPX6 protein using CODEML, branch model = 2. The P-values for significance in the difference between the likelihood of the branch model to the null model for each domain are shown in Table 1. Calculations of dN/dS ratios over protein domains were repeated for the other genes in the GPX family (Table 1).

### dN/dS Ratios in GPX3 (sloth and kangaroo rat)

We found that the Sec-containing GPX3 protein lacks its defining Sec residue in two species; the Hoffman’s two-toed sloth and the kangaroo rat. Here, the Sec has been exchanged for either Glutamine (in the case of the sloth) or for Serine (in the case of the kangaroo rat). Because of these exchanges, both the sloth-GPX3 protein and kangaroo rat-GPX3 protein were excluded from the branch model analysis of dN/dS in GPX3 (see “dN/dS ratios in GPX proteins”). However, these proteins were then included (and following alignment and HMMER analysis steps repeated) to allow Branch-site analysis on these two branches (see “Inferring Selection on the GPX6 Sites”; Table S1).

### Inferring Selection on the GPX6 Sites

To test for selection acting on individual sites across the entire tree, we employed the Site model in PAML (Yang, 2007). We compared model 7 (beta; model = 0, NSsites=7) to model 8 (beta plus selection; model =0, NSsites=8). Here, model 7 is the null model of a beta distributed variable selective pressure across sites, whereas model 8 is the beta distributed model plus selection.

Given the significance of this site model test, we then tested for selection acting on sites in the GPX domain along particular branches across the tree using the Branch-site model in PAML. This model (model = 2, NSsites=2) allows the dN/dS ratio to vary both amongst the sites and amongst the specified foreground and background branches and outputs the probability of each site being under selection in the foreground branches according to PAML’s Bayes Empirical Bayes (BEB) inference method (Yang et al., 2005).

This model classifies the sites into those that have dN/dS values that remain the same on the foreground and background branches (*ω* < 1 or *ω* = 1 in both branches) and those that differ amongst the branches (*ω* < 1 or *ω* = 1 in background branches and *ω* > 1 in foreground branches), outputting the proportion of each site class. This method then calculates the posterior probability of each site being under selection in the foreground branches, whilst accounting for sampling errors by using a Bayesian prior (Yang et al., 2005). This model is compared to the corresponding null model, which is the same in all ways apart from the fixation of *ω*_2_.

We used this model initially on the branches where Sec was inferred to be lost (Fig 1; dashed green branches acting as the foreground) and found significant evidence for selection on particular sites within this region (Table S2). We then extended this model to test along the entire protein region for these same branches. However, given the non-significant result, we repeated the use of this model to the closest related lineages where Sec was inferred to be lost (see Table S2). This is because this model is suited to identifying a set of sites under selection across the entire set of foreground branches; if selection is acting across different sites in the foreground branches, as somewhat expected given the role of epistasis over more diverged lineages, this model will return a non-significant result. That is to say, we expect epistasis to place a limit the number of the same sites being under selection across more diverged lineages.

Given we see significant evidence for selection across the mostly closer related lineages where Sec was inferred to be lost (the branch leading to squirrel monkey-marmoset, the Eumuroida branch and the branch leading to rabbit; see Fig 1), we are able to test if these probabilities are enriched in certain subsets of sites using Mann-Whitney U tests (see “Identifying Convergent Changes across Cys-Branches” and “Convergence in the *Eumuroida* Lineage”).

### Identifying Convergent Changes across Cys-Branches

Convergent changes in GPX6 across lineages were identified using CONVERG2 (Zhang & Kumar, 1997) This programme differentiates between parallel and convergent amino acid changes (parallel changes being those changes from the same ancestral amino acid to the same derived amino acid, convergent changes being those changes from a different ancestral amino acid to the same derived amino acid) but for brevity, both are referred to as convergent amino acid changes here.

Convergent changes were identified between the GPX6_Cys_ lineages; either the branches where Sec was exchanged for Cys or the species branches where Cys was maintained (the dashed and solid green lines in Fig 1, respectively). The observed frequency of these convergent changes was then compared with the expected frequency of convergent changes, also calculated using CONVERG2.

Since the pathway to recover catalytic activity may not be limited to the same amino acid changes, but may be restricted to particular sites in the protein, the CONVERG2 programme was also edited to identify convergent site changes, which do not condition on resulting in the same amino acid across branches. Hence, convergent site changes are used to identify sites that show repeated changes across lineages. Both convergent amino acid and site changes are referred to as convergent in the paper, but the exact nature of each convergent change can be seen in Tables S3-9.

Where the sequences for the species containing Cys in GPX6 were available, the equivalent analyses were run on the Sec-containing GPX proteins (GPX1, 2, 3 and 4) and the Cys-containing proteins (GPX7 and 8). We advise focusing on the CONVERG2 results for GPX3 and GPX5 for two reasons: 1) these are the immediate paralogues to GPX6; and 2) the gaps in the other proteins do not allow, we believe, a full representation of the potential instances of convergence. Still, we do stress that the pattern over all of these non-GPX6 proteins is that of much reduced convergence, if any, across the Cys-lineages identified in GPX6.

Using the posterior probabilities of selection obtained from the BEB results of the Branch-site model when the foreground branches are specified as the branch leading to squirrel monkey-marmoset, the Eumuroida branch and the branch leading to rabbit (*i.e*., using the probabilities of sites being under selection in these three branches), we also compared the posterior probability of selection on convergent sites. Here, we exclude convergent sites only identified using either the cat or walrus terminal branches, since they are excluded from the probability calculation. To test for enrichment, we use a Mann-Whitney U test to account for the nonparametric data.

### Convergence in the *Eumuroida* Lineage

By inferring the GPX6 protein sequences along the *Eumuroida* branch (see “Inferring the Loss of Sec in GPX6”) we identified 26 sites that changed over the *Eumuroida* branch. Using the results from CONVERG2, we were then able to identify 15 sites that show signatures of convergence across GPX6_Cys_ lineages (to the exclusion of those identified from cat and walrus) and show the distribution within each protein domain. We were also able to infer a further 22 amino acid sites (19 substitutions and a 3 C-terminal extension) that changed between the end of the *Eumuroida* branch (dashed green branch in Figure 2a) and the modern mouse GPX6 protein; m-GPX6_Cys+22_. CONVERG2 was again used to identify which of these 19 substitutions signatures of convergence across GPX6_Cys_ lineages.

These subsets of sites, both the 15 along the *Eumuroida* branch and the 8 leading to modern day mouse, were tested for enrichment of selection signatures by comparing the posterior probabilities of selection using a Mann-Whitney U test. Here, we again use these posterior probabilities are the probabilities of sites being under selection in the branch leading to squirrel monkey-marmoset, the Eumuroida branch and the branch leading to rabbit.

Using PHYML, we reconstructed the mammalian tree given: a) the full GPX6 gene, b) the N-terminal of GPX6, c) the GPX domain of GPX6, d) the N-terminal of GPX6, e) the 26 sites that change across the *Eumuroida* branch (Fig S3) as well as the 14 sites that show changes across the *Eumuroida* branch and convergent changes across *GPx*6_*Cys*_ branches (Fig 2b). This was repeated using the full GPX3 and GPX5 proteins (Fig S4).

### Expected Levels of Convergence given rate of amino acid exchange

The evolution of the GPX6 protein sequence across our mammalian phylogeny was simulated using Seq-Gen (Rambaut & Grassly, 1997). This simulated evolution begins with the inferred ancestral sequence at the base of our mammalian clade (see “Inferring the Loss of Sec in GPX6”) and extends to all modern mammalian proteins, using the JTT model of amino acid substitution. Tree lengths were given by the rate of amino acid changes along each branch of the mammalian tree as from the calculated dN value in the CODEML package from PAML((Yang, 2007); see “dN/dS ratios in GPX proteins”). Hence, this simulation recreates chance amino acid exchanges along each branch at its observed rate.

Having run the simulation 1,000 times, we then use CONVERG2 (Zhang & Kumar, 1997) to identify convergent site changes between the lineages where Sec was lost for Cys (solid green lines in Fig 1) for each simulation run (see “Identifying Convergent Changes across Cys-Branches”). The distribution of convergent changes under this expected rate of amino acid exchange is then plotted, and compared to the observed number of convergent site changes (Fig S5). The equivalent simulations were run for all other GPX proteins, and we further compared the observed and expected number of convergent site changes for these proteins (Figs S7-13)

To confirm that the higher number of observed convergent changes relative to our expectation are focused within the functional GPX domain, and this isn’t simply an artefact of elevated evolutionary rate within this domain, we also repeated these simulations on only the GPX domain. Here, the tree lengths were given by the rate of amino acid changes (dN) from the GPX domain only. Comparisons between observed and expected convergent site changes are given in Fig S6.

### Ancestral Reconstruction of GPX6 along the *Eumuroida* Lineage

Following our inference of ancestral sequences (see “Inferring the Loss of Sec in GPX6”), we were able to reconstruct 3 ancient proteins along the *Eumuroida* lineage. These proteins are: 1) the protein just prior to the loss of Sec in the ancestor of *Eumuroida* (Eu-GPX6_Sec_); 2) the same ancestral protein but with Sec exchanged for Cys (Eu-GPX6_Cys_); 3) the protein at the derived end of the *Eumuroida* branch, now containing the additional 25 sites that have changed along the *Eumuroida* branch (Eu-GPX6_Cys+25_).

As previously described (see “Inferring the Loss of Sec in GPX6”), the residue with most support from the four inferred sequences was taken as the consensus residue for each site, with the exception of site 54 in Eu-GPX6_Cys+25_. Here, the consensus residue was taken as “Q” despite the methods used suggesting “H” since “H” is not present at that site for any of the present-day species. Of the 217 amino acid sites, 208 (95.85%) were resolved unanimously across the four inference methods. Of the remaining 9 sites (4.15%) that were inferred differently across them methods, 7 (3.23% of total sites) of these sites differed across the inference of the Eu-GPX6_Sec_ protein and 2 (0.92% of total sites) differed across the inference of the Eu-GPX6_Cys+25_.

These consensus sequences provide the final Eu-GPX6_Sec_ and Eu-GPX6_Cys+25_ proteins. As before, sites with an average posterior probability below 0.9, as calculated using HMMER (Potter et al., 2018) were removed from subsequent PAML analysis.

### Experimental Reconstruction and Enzymatic Assay of Ancient GPX6 Proteins

We reconstructed the Eu-GPX6_Sec_, Eu-GPX6_Cys_ and Eu-GPX6_Cys+25_ proteins from heterologous expression in *Escherichia coli*. We used a mutant *E.coli* strain that does not recognise UAG as a STOP codon, resulting in a much higher yield of Eu-GPX6_Sec_ that would otherwise be produced by *E.coli* with standard genetic code decoding (Cheng & Arnér, 2017). The catalytic activity of each protein was evaluated by measuring the peroxidation activity on H_2_O_2_. This was used to compare activity between Eu-GPX6_Sec_ and Eu-GPX6_Cys_ (Fig 3), but we were not able to reliably measure peroxidation activity for Eu-GPX6_Cys+25_ owing to limited yield of this protein, stemming from its poor solubility.The catalytic activity of each protein was evaluated by measuring the peroxidation activity on H_2_O_2_. This was used to compare activity between Eu-GPX6_Sec_ and Eu-GPX6_Cys_ (Fig 3), but we were not able to reliably measure peroxidation activity for Eu-GPX6_Cys+25_ owing to limited yield of this protein, stemming from its poor solubility.

### Molecular docking simulations

Structures for the GPX6 orthologs and nodes of the ancestral sequence reconstructions were built using AlphaFold2 (Jumper et al., 2021). All protein sequences considered cysteines at their catalytic positions, given the inability to represent non-canonical residues for the *ab initio* model construction. We ran protein-ligand binding energy landscape explorations using the PELE software (Borrelli et al., 2005) for each protein structure. Ligands for the simulation were glutathione and glutathione disulfide. Simulations were first run to discover catalytic poses with low global energies; the catalytic distance was considered as the closest sulphur-sulphur distance between the catalytic cysteine and the glutathione sulphur atoms. The lowest binding energy poses, filtered by a catalytic distance threshold below 4Å, were used to run a second PELE simulation, thus focusing on exploring this catalytic minimum binding energy configuration. Each simulation comprised 95 replicas of 100 equilibration steps that constrained the ligand to its starting position, followed by 1000 PELE steps without any constraint over the ligand coordinates.

All simulation trajectories for the same ligand were simultaneously analyzed using as features all ligand positions aligned to a common protein reference structure. A Time-structure Independent Component Analysis (TICA) was built to find the common slowest-relaxing feature combination (Molgedey & Schuster, 1994) with the PyEMMA library (Scherer et al., 2015). Finally, and separately for each protein and ligand simulation, the probabilities of visiting the slowest TICA coordinate (IC1) according to the catalytic distance (S-S) were plotted as a free energy map (Fig 4 and Fig S14).

## Acknowledgements

EA and QC funded by the Karolinska Institutet, The Knut and Alice Wallenberg Foundations (KAW 2019.0059), The Swedish Cancer Society (21 1463 Pj), The Swedish Research Council (2021-02214), National Laboratories Excellence program under the National Tumor Biology Laboratory project (2022-2.1.1-NL-2022-00010) and the Hungarian Thematic Excellence Programme (TKP2021-EGA-44) and The National Research, Development and Innovation Office (NKFIH) grant ED_18-1-2019-0025. MF, BOM, JVF are funded by project BIO2017-83650-P, financed by Spanish Ministerio de Ciencia e Innovación (MCIN). A.A. is funded by UCL’s Wellcome Institutional Strategic Support Fund 3 (grant reference 204841/Z/16/Z). S.C. and J.R. are funded by NIHR GOSH BRC. The views expressed are those of the authors and not necessarily reflect those of the funding body, including those of the NHS, the NIHR, or the Department of Health. Figures 1,2, S2, S5-S13 created with the use of Biorender.

