## Supplemental Information for "Ancient loss of catalytic selenocysteine spurred convergent adaptation in a mammalian oxidoreductase"

#### Abandoning selenocysteine for cysteine reappraises enzyme catalysis

Jasmin Rees[1,2*], Gaurab Sarangi[3*], Qing Cheng[4*], Martin Floor[5,6*], Aida M Andrés[2], Baldomero Oliva Miguel[7], Jordi Villà-Freixa[5,8], Elias SJ Arnér[4,9], Sergi Castellano[1,10]

[1] Great Ormond Street Institute of Child Health, University College London, London, United Kingdom
[2] Division of Biosciences, University College London, London, United Kingdom
[3] Department of Evolutionary Genetics, Max Planck Institute for Evolutionary Anthropology, Leipzig, Germany
[4] Division of Biochemistry, Department of Medical Biochemistry and Biophysics, Karolinska Institutet, Stockholm, Sweden
[5] Department of Biosciences, Faculty of Sciences and Technology, Universitat de Vic - Universitat Central de Catalunya, Vic, Spain
[6] Barcelona Supercomputing Center (BSC), Barcelona, Spain
[7] Department of Health and Experimental Sciences, Universitat Pompeu Fabra, Barcelona, Spain
[8] Institut de Recerca i Innovació en Ciències de la Vida i de la Salut a la Catalunya Central (IRIS-CC), Vic, Spain
[9] Department of Selenoprotein Research, National Institute of Oncology, Budapest, Hungary
[10] UCL Genomics, University College London, London, United Kingdom

[*] These authors contributed equally.

### Figures

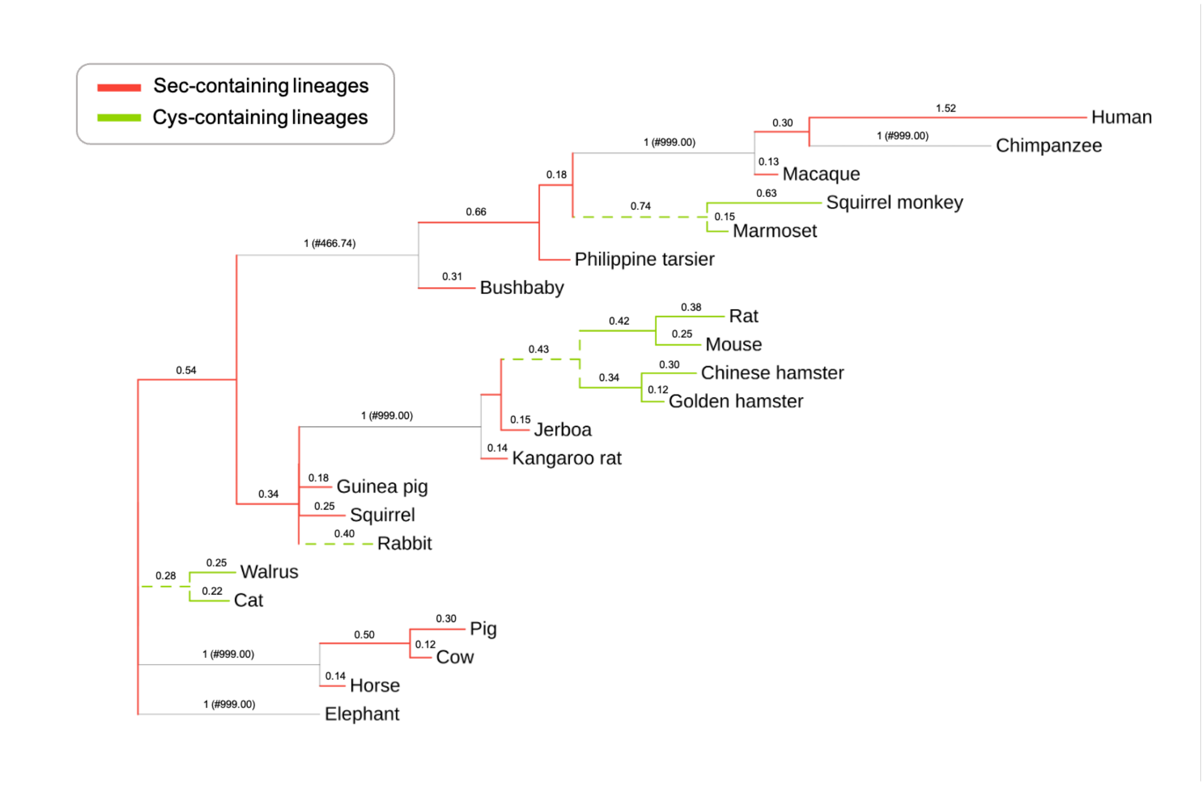


**Figure S1:** Phylogeny of the 22 mammals in our analysis. In red, $GPX6_{Sec}$ branches, in green, $GPX6_{Cys}$ ones. Branch lengths are proportional to their corresponding dN/dS as estimated by the free-ratio model in PAML. In some lineages, PAML estimates very few synonymous changes compared to non-synonymous changes and this results in an unnaturally large dN/dS value. These branches are assigned a dN/dS ratio of 1 and coloured grey, with the actual ratio estimated by PAML is given in parenthesis (#). Branches with dN/dS values given as less than 0.01 are not labelled.

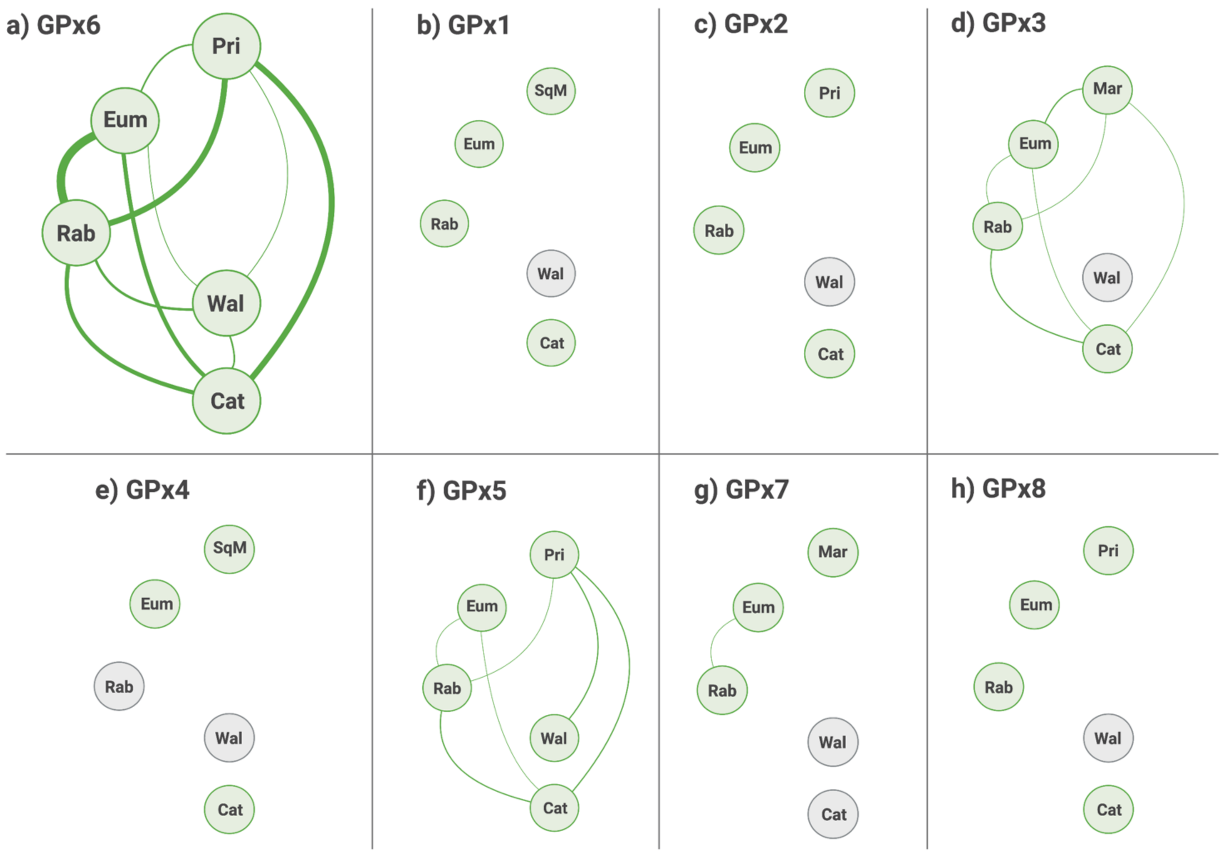


**Figure S2.** Schematic diagram demonstrating the convergence between branches in GPX6 where Sec was inferred to have been lost. Connection thickness is proportional to the number of convergent sites identified. When a species protein is unavailable, the node is in grey. Pri = primate branch (leading to squirrel monkey and marmoset; Fig 1); Eum= Eumuroida;  Rab = Rabbit; Wal = Walrus; Cat = Cat; SqM = Squirrel monkey; Mar = Marmoset

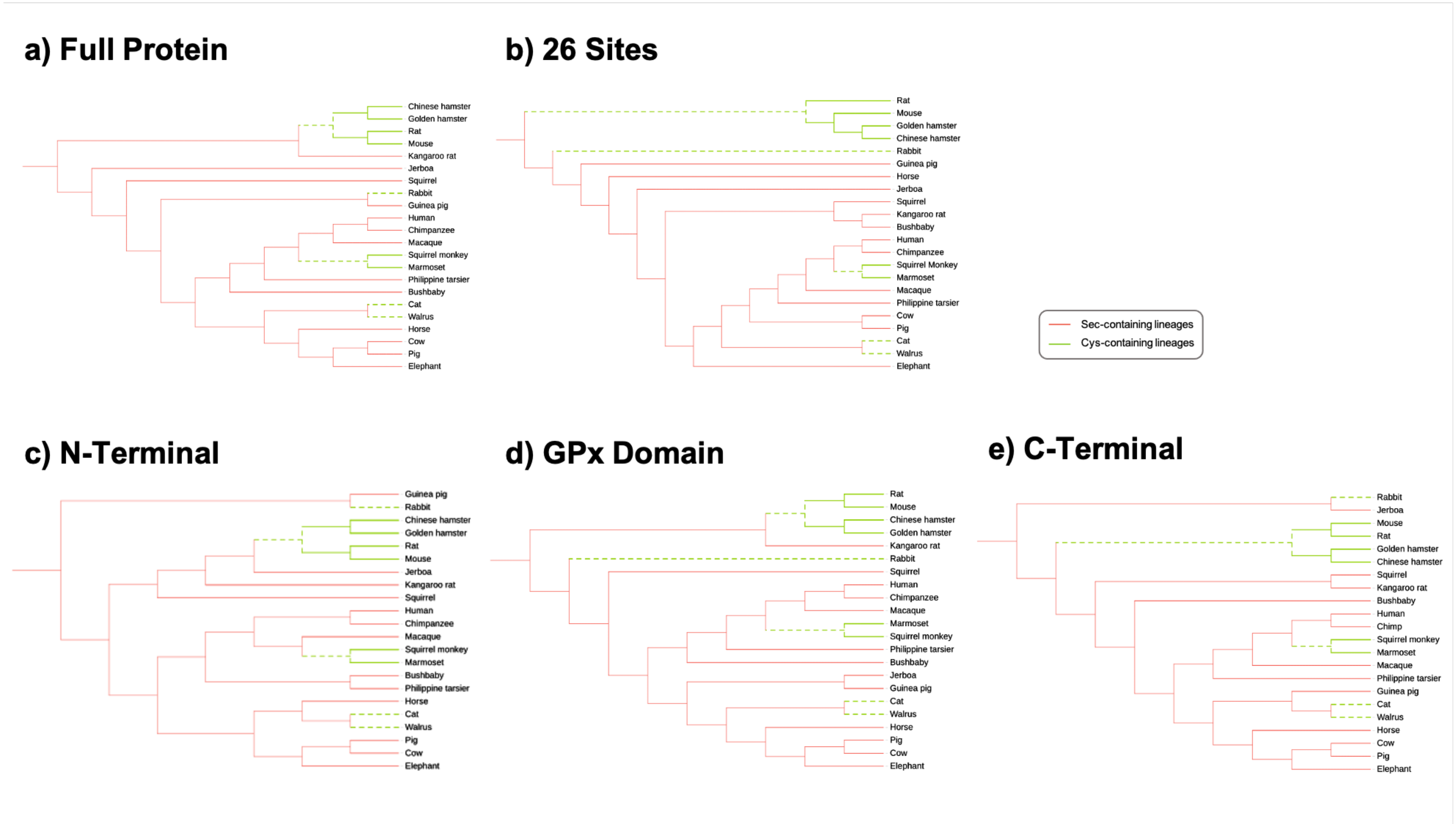


**Figure S3.** Topology of the phylogenetic tree for GPX6, with midpoint rooting, constructed using PHYML from the **a**) full GPX6 protein; **b**) the 26 sites that differ between Eu-GPX6$_{Sec}$ and Eu-GPX6$_{Cys+25}$; **c**) the N-terminal of GPX6; **d**) the GPX domain of GPX6 and **e**) the C-terminal of GPX6. In red, GPX6$_{Sec}$ branches, in green, GPX6$_{Cys}$ ones. Dashed green branches represent GPX6$_{Cys}$ lineages at the time Sec was lost.

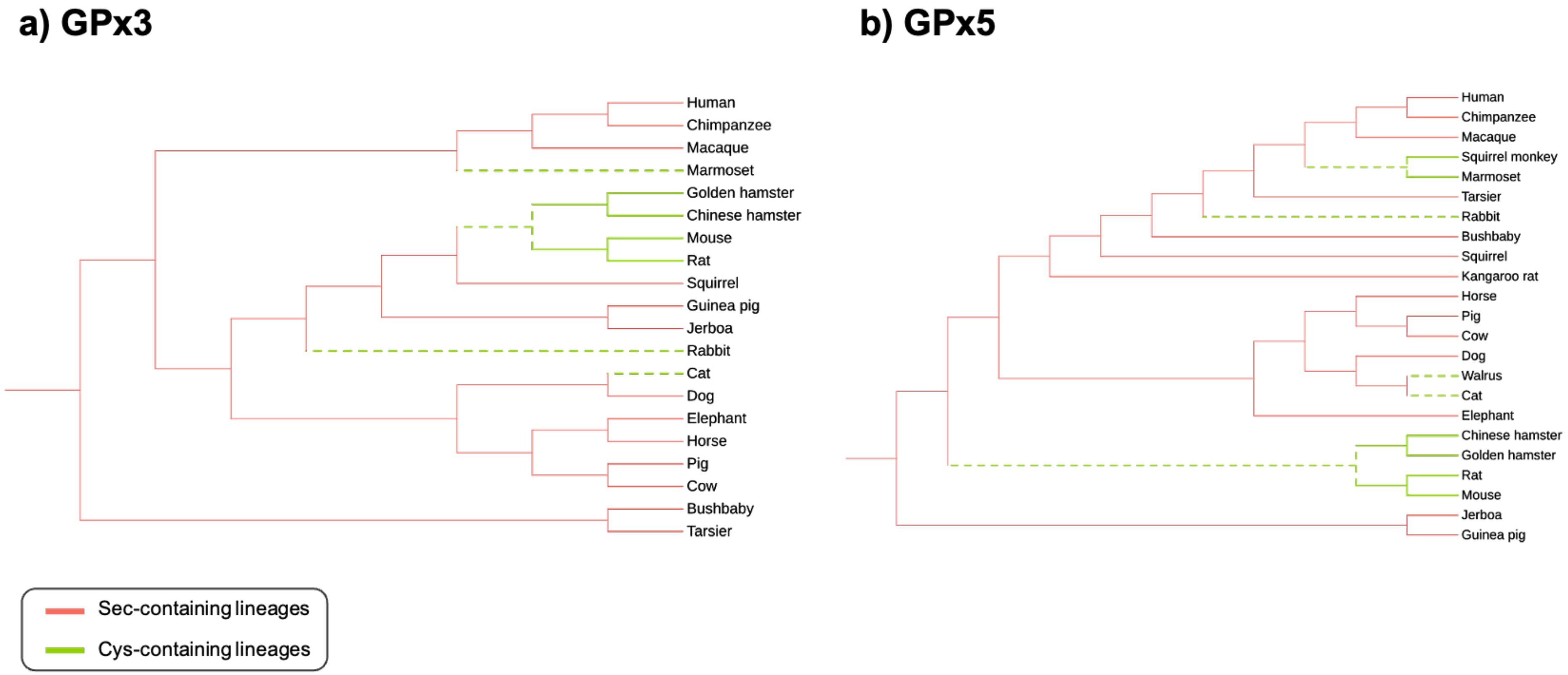


**Figure S4.** Topology of the phylogenetic trees, with midpoint rooting, constructed using PHYML from the available mammalian proteins of a) GPX3 and b) GPX5. In green, $GPX6_{Cys}$ ones. Dashed green branches represent $GPX6_{Cys}$ lineages at the time Sec was lost. The species names in italics indicate those species not represented in our wider GPX6 analysis.

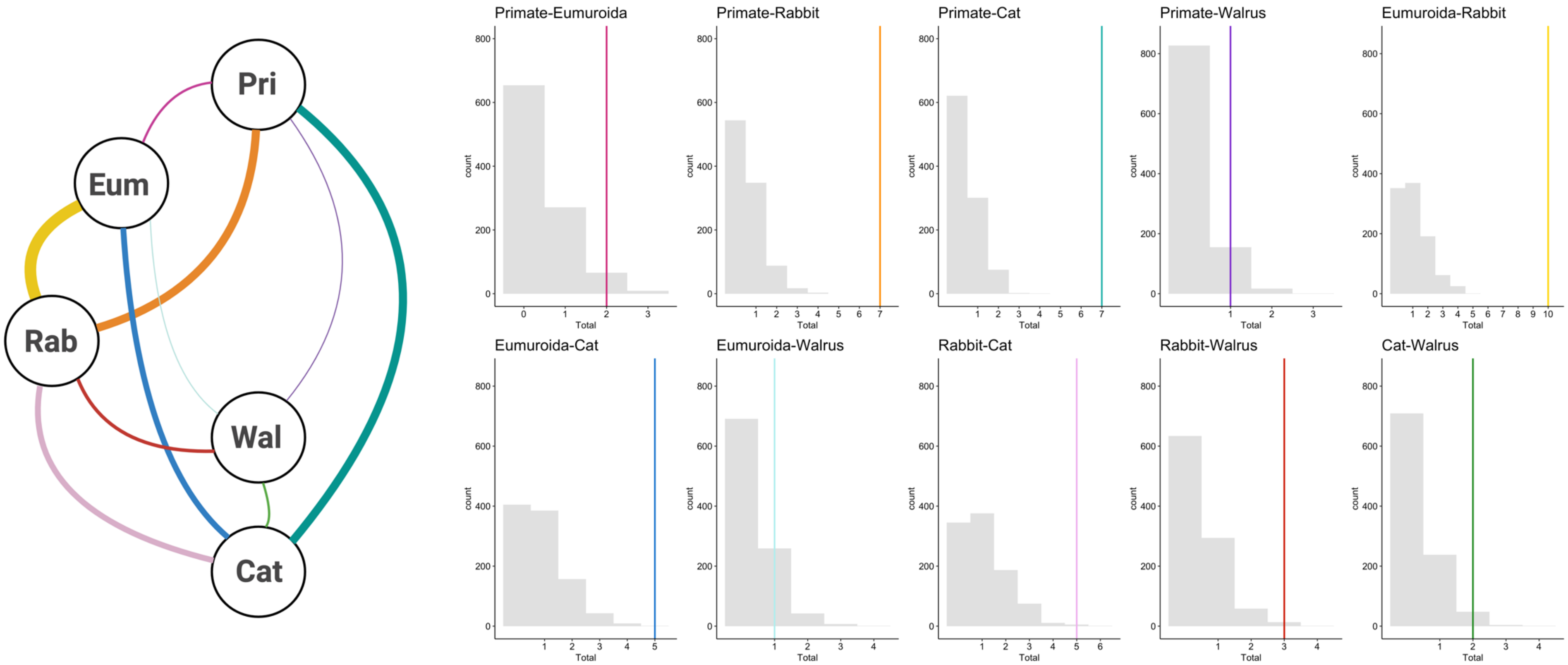


**Figure S5.** Schematic representation of the number of observed convergence in the GPX6 protein between lineages where Sec is lost for Cys, where thickness of the line represents the number of convergent changes (left). Expected distribution of convergent changes in the full GPX6 protein between lineages where Sec is lost for Cys according to our Seq-Gen simulations, where the observed numbers of convergent changes are given by coloured lines (right).

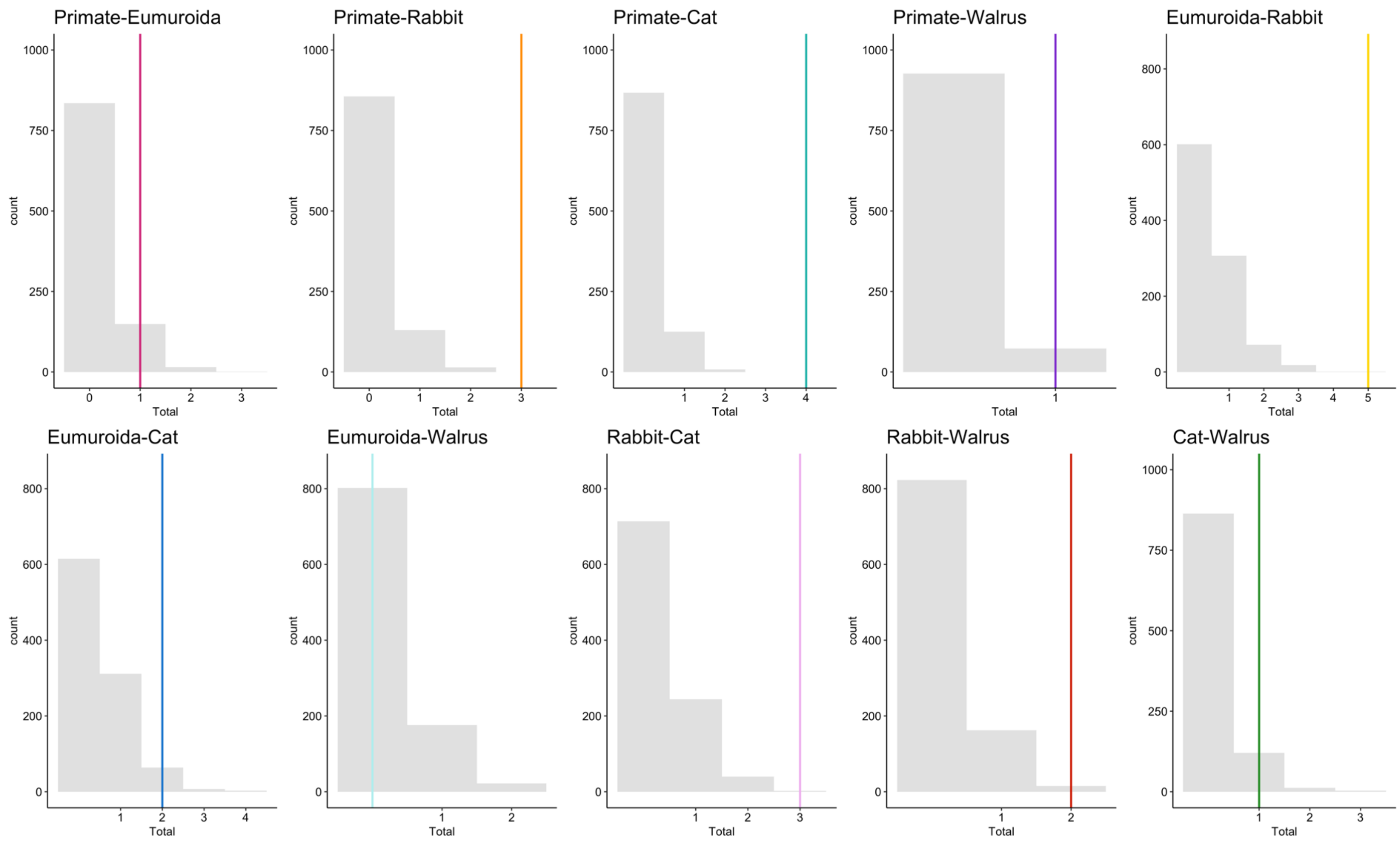


**Figure S6.** Expected distribution of convergent changes in the GPX domain of the GPX6 between lineages where Sec is lost for Cys according to our Seq-Gen simulations, where the observed numbers of convergent changes are given by coloured lines.

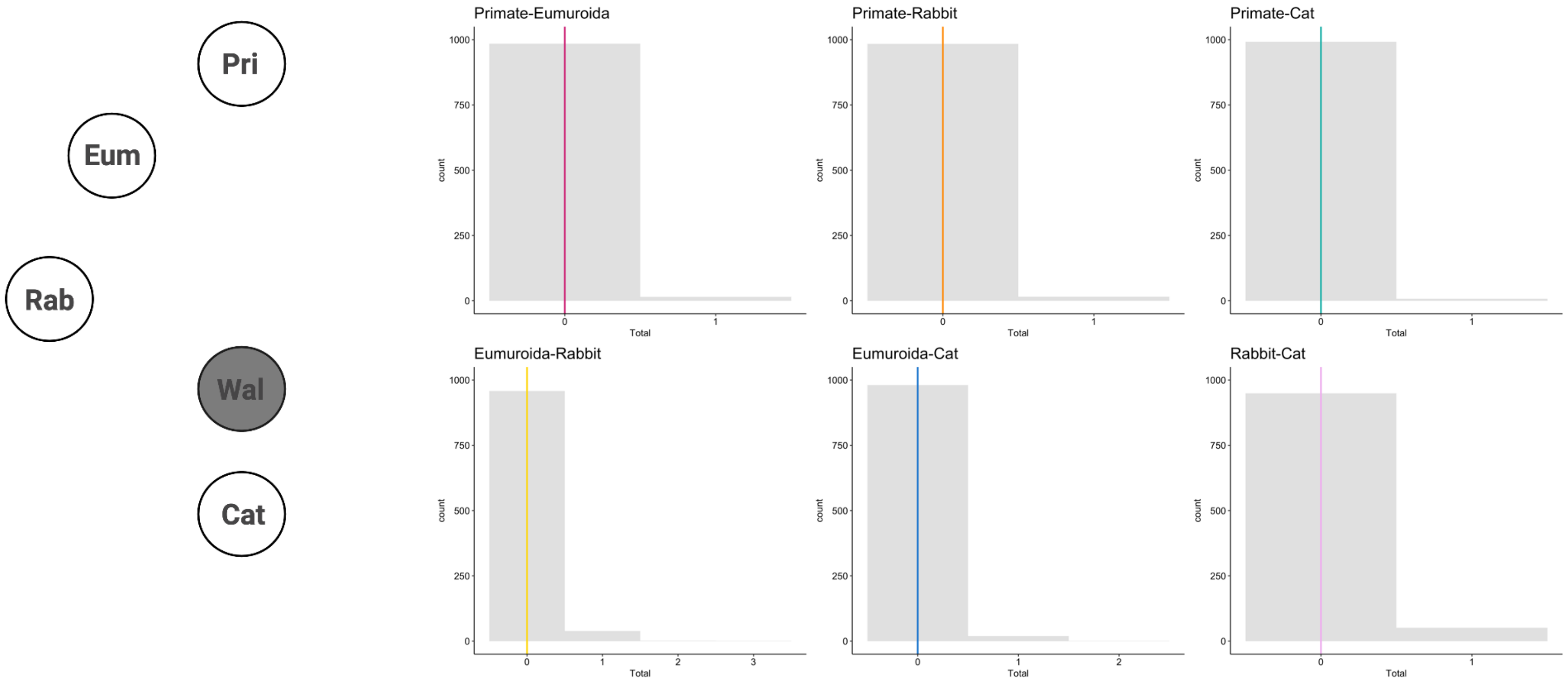


**Figure S7.** Schematic representation of the number of observed convergence in the GPX1 protein between lineages where Sec is lost for Cys in GPX6, where thickness of the line represents the number of convergent changes (left). Expected distribution of convergent changes in GPX1 between lineages where Sec is lost for Cys in GPX6 according to our Seq-Gen simulations, where the observed numbers of convergent changes are given by coloured lines (right).

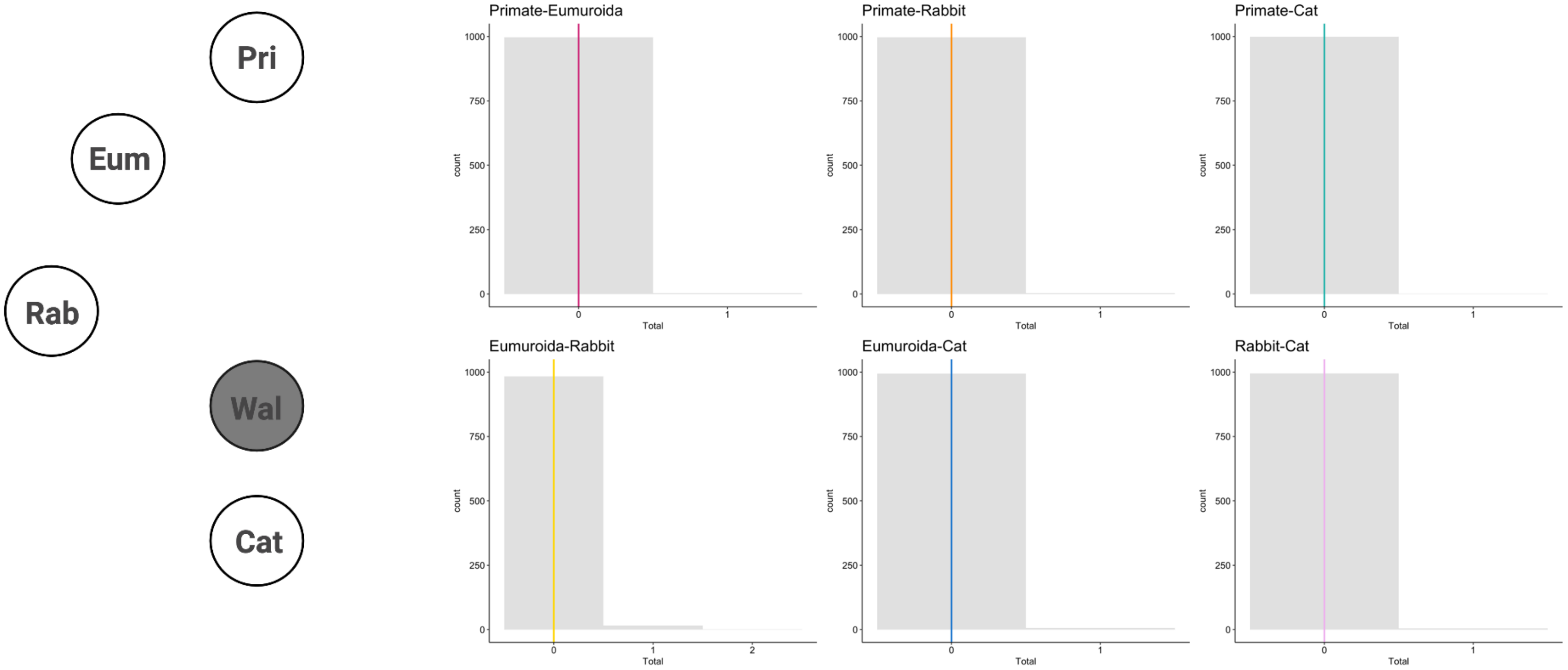


**Figure S8.** Schematic representation of the number of observed convergence in the GPX2 protein between lineages where Sec is lost for Cys in GPX6, where thickness of the line represents the number of convergent changes (left). Expected distribution of convergent changes in GPX2 between lineages where Sec is lost for Cys in GPX6 according to our Seq-Gen simulations, where the observed numbers of convergent changes are given by coloured lines (right).

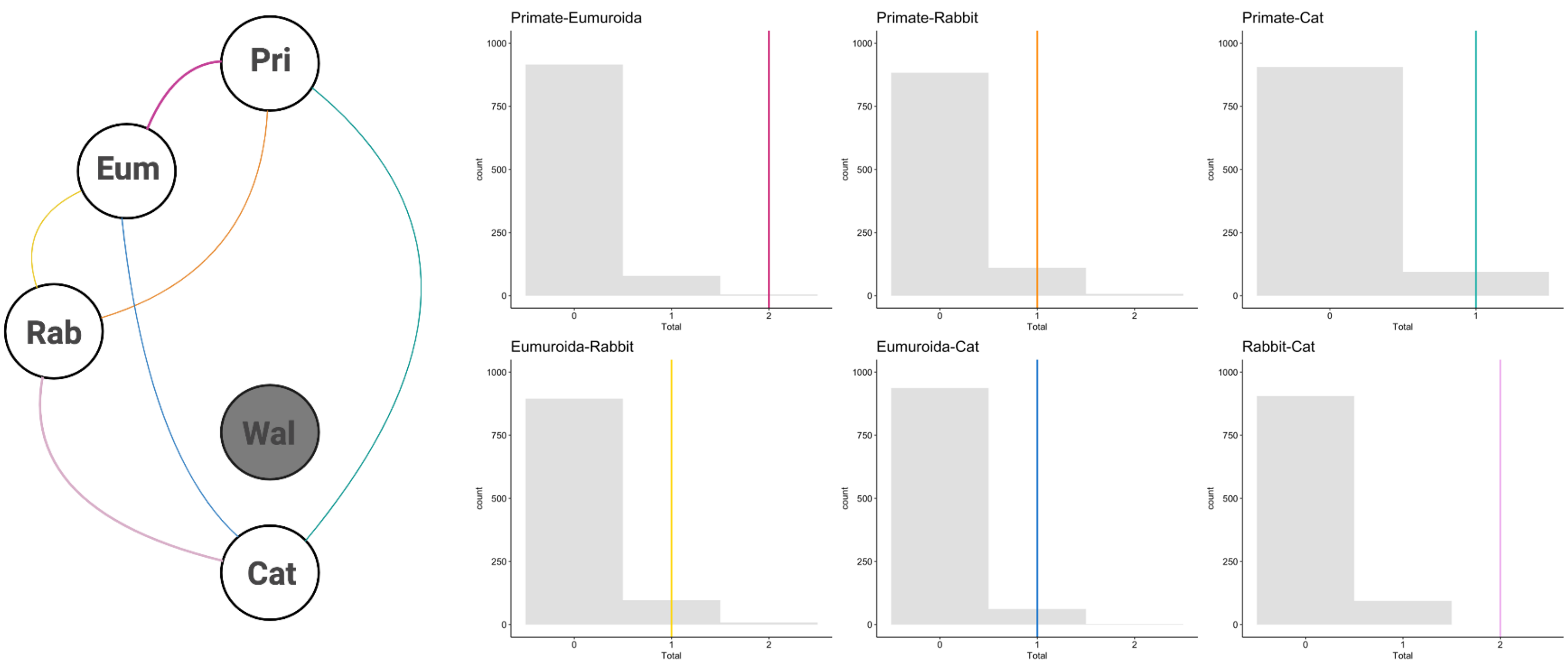


**Figure S9.** Schematic representation of the number of observed convergence in the GPX3 protein between lineages where Sec is lost for Cys in GPX6, where thickness of the line represents the number of convergent changes (left). Expected distribution of convergent changes in GPX3 between lineages where Sec is lost for Cys in GPX6 according to our Seq-Gen simulations, where the observed numbers of convergent changes are given by coloured lines (right).

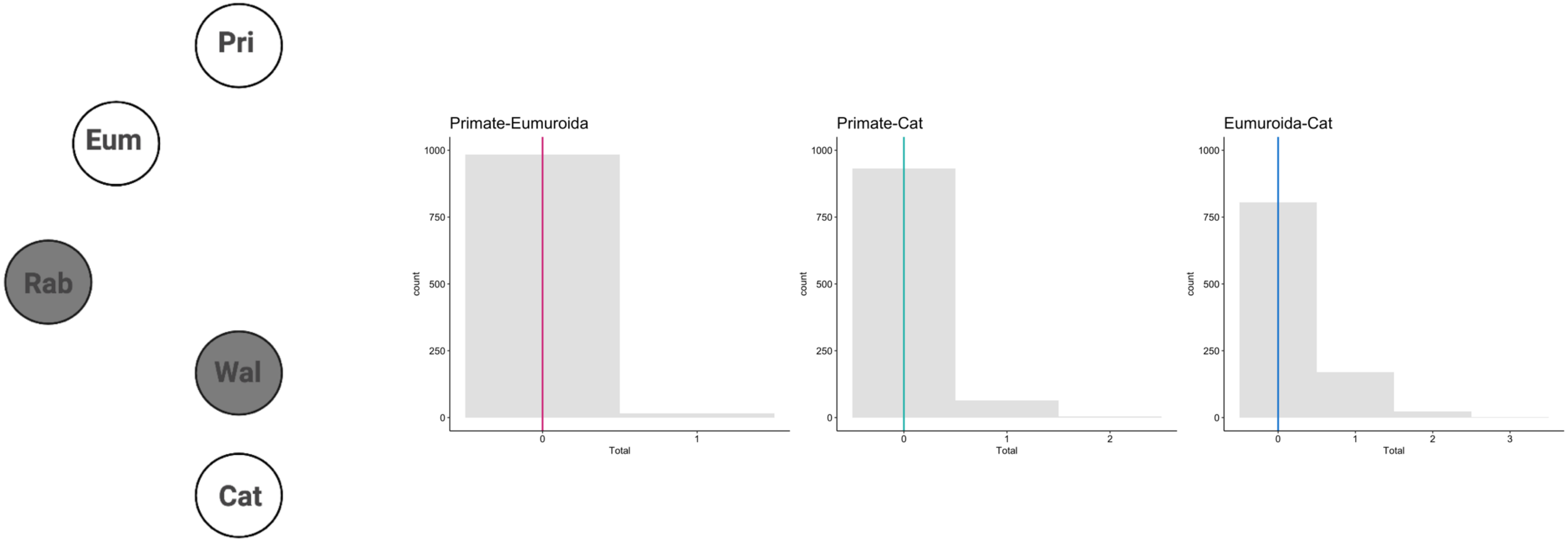


**Figure S10.** Schematic representation of the number of observed convergence in the GPX4 protein between lineages where Sec is lost for Cys in GPX6, where thickness of the line represents the number of convergent changes (left). Expected distribution of convergent changes in GPX4 between lineages where Sec is lost for Cys in GPX6 according to our Seq-Gen simulations, where the observed numbers of convergent changes are given by coloured lines (right).

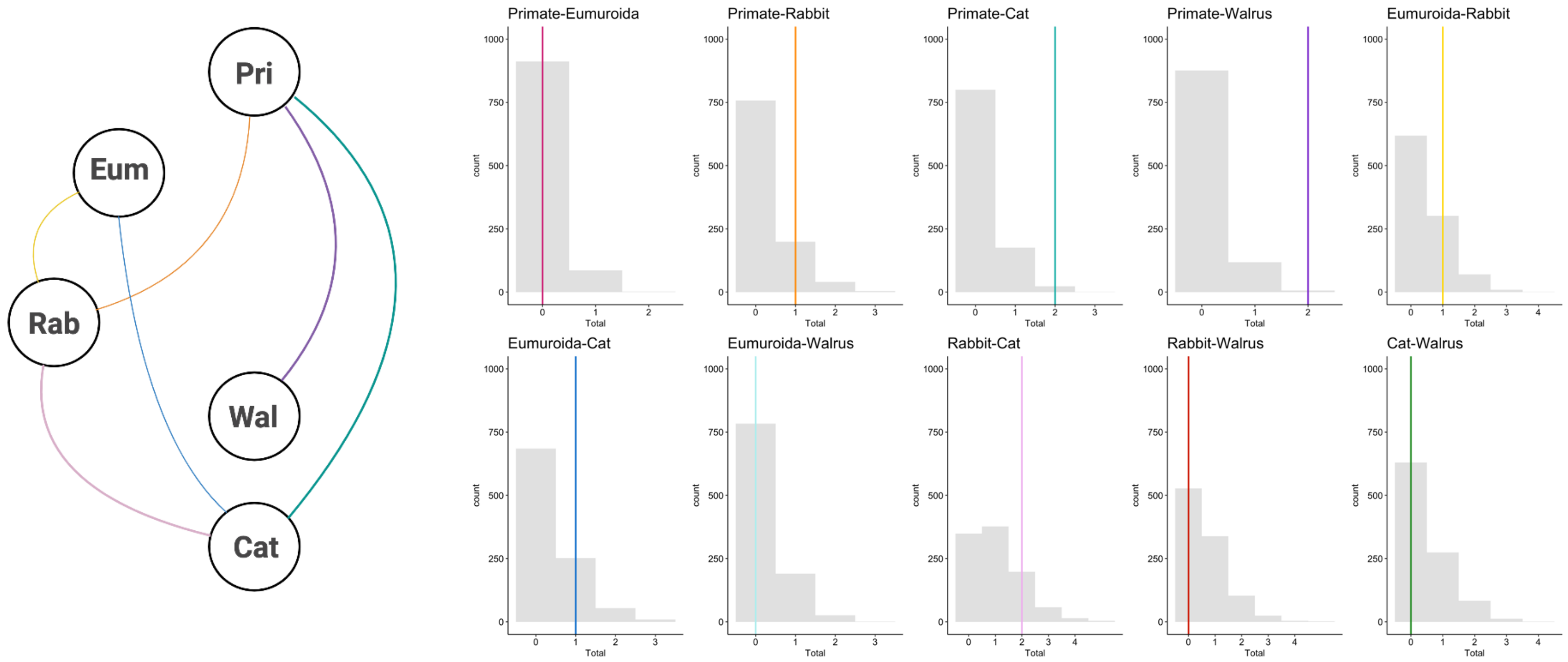


**Figure S11.** Schematic representation of the number of observed convergence in the GPX5 protein between lineages where Sec is lost for Cys in GPX6, where thickness of the line represents the number of convergent changes (left). Expected distribution of convergent changes in GPX5 between lineages where Sec is lost for Cys in GPX6 according to our Seq-Gen simulations, where the observed numbers of convergent changes are given by coloured lines (right).

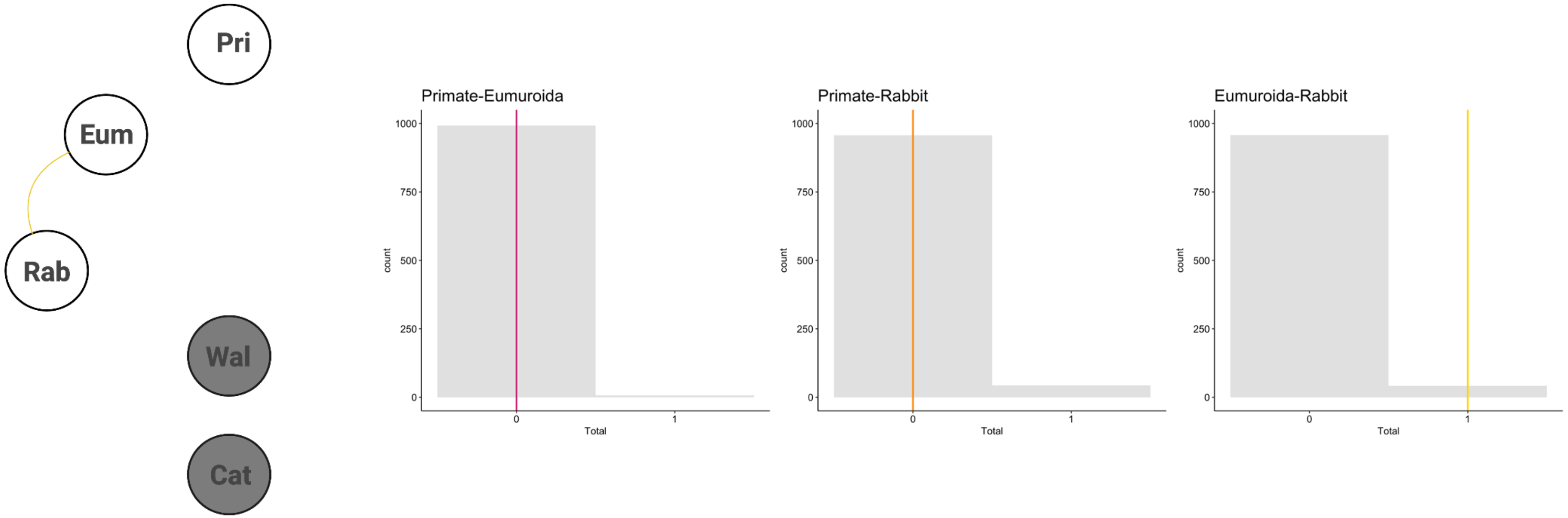


**Figure S12.** Schematic representation of the number of observed convergence in the GPX7 protein between lineages where Sec is lost for Cys in GPX6, where thickness of the line represents the number of convergent changes (left). Expected distribution of convergent changes in GPX7 between lineages where Sec is lost for Cys in GPX6 according to our Seq-Gen simulations, where the observed numbers of convergent changes are given by coloured lines (right).

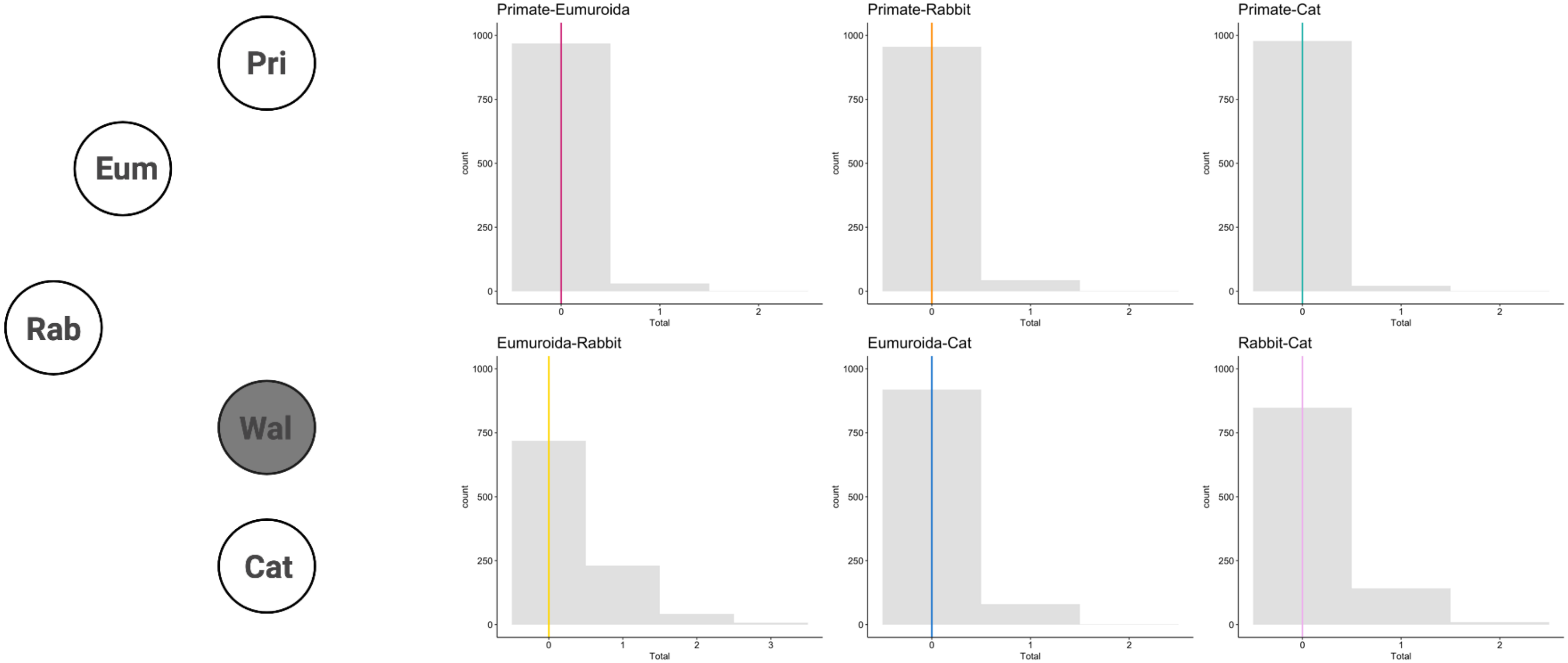


**Figure S13.** Schematic representation of the number of observed convergence in the GPX8 protein between lineages where Sec is lost for Cys in GPX6, where thickness of the line represents the number of convergent changes (left). Expected distribution of convergent changes in GPX8 between lineages where Sec is lost for Cys in GPX6 according to our Seq-Gen simulations, where the observed numbers of convergent changes are given by coloured lines (right).

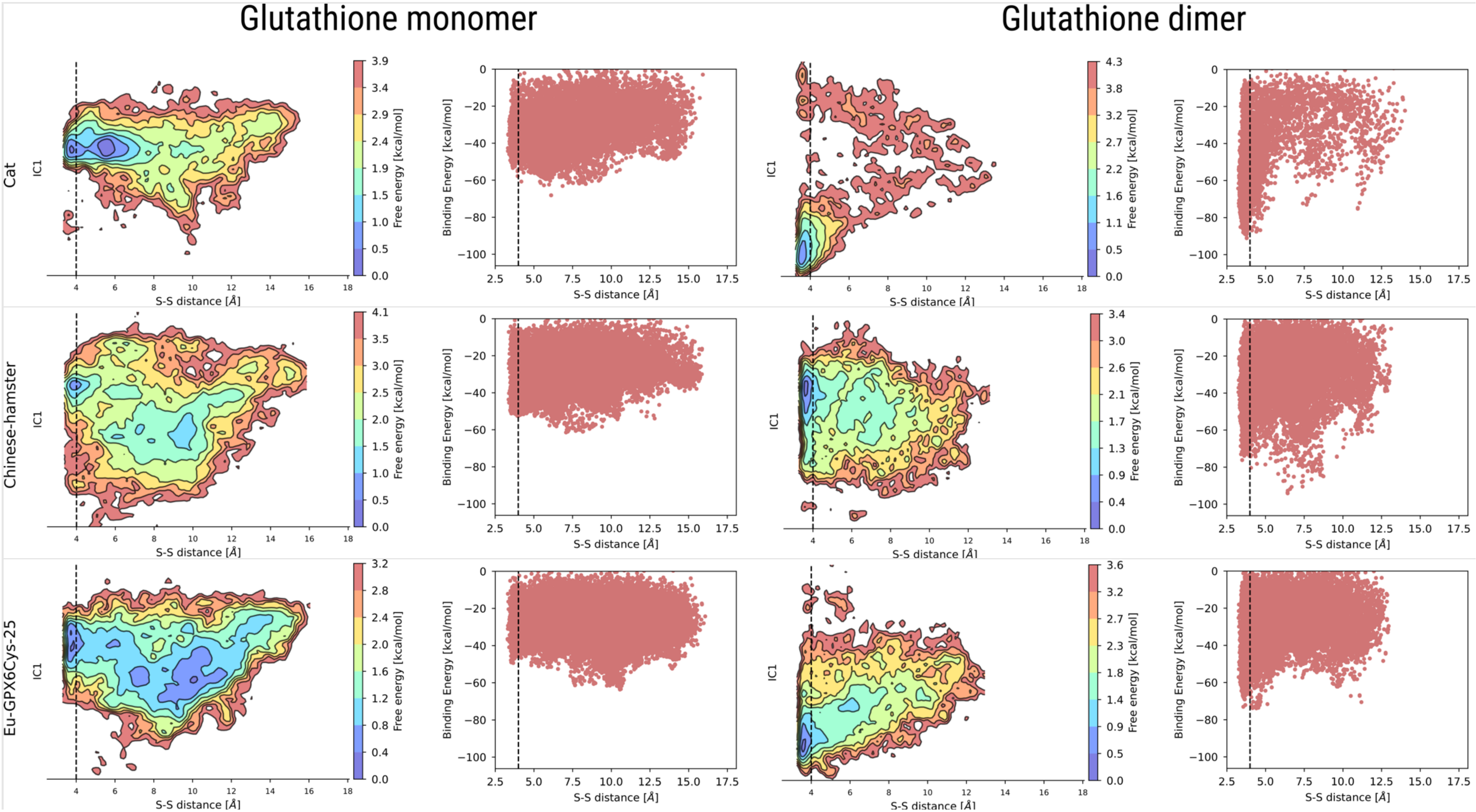
Glutathione monomer
Glutathione dimer
Cat
Chinese-hamster
Eu-GPX6Cys-25
IC1
S-S distance [Å]
Free energy [kcal/mol]
Binding Energy [kcal/mol]

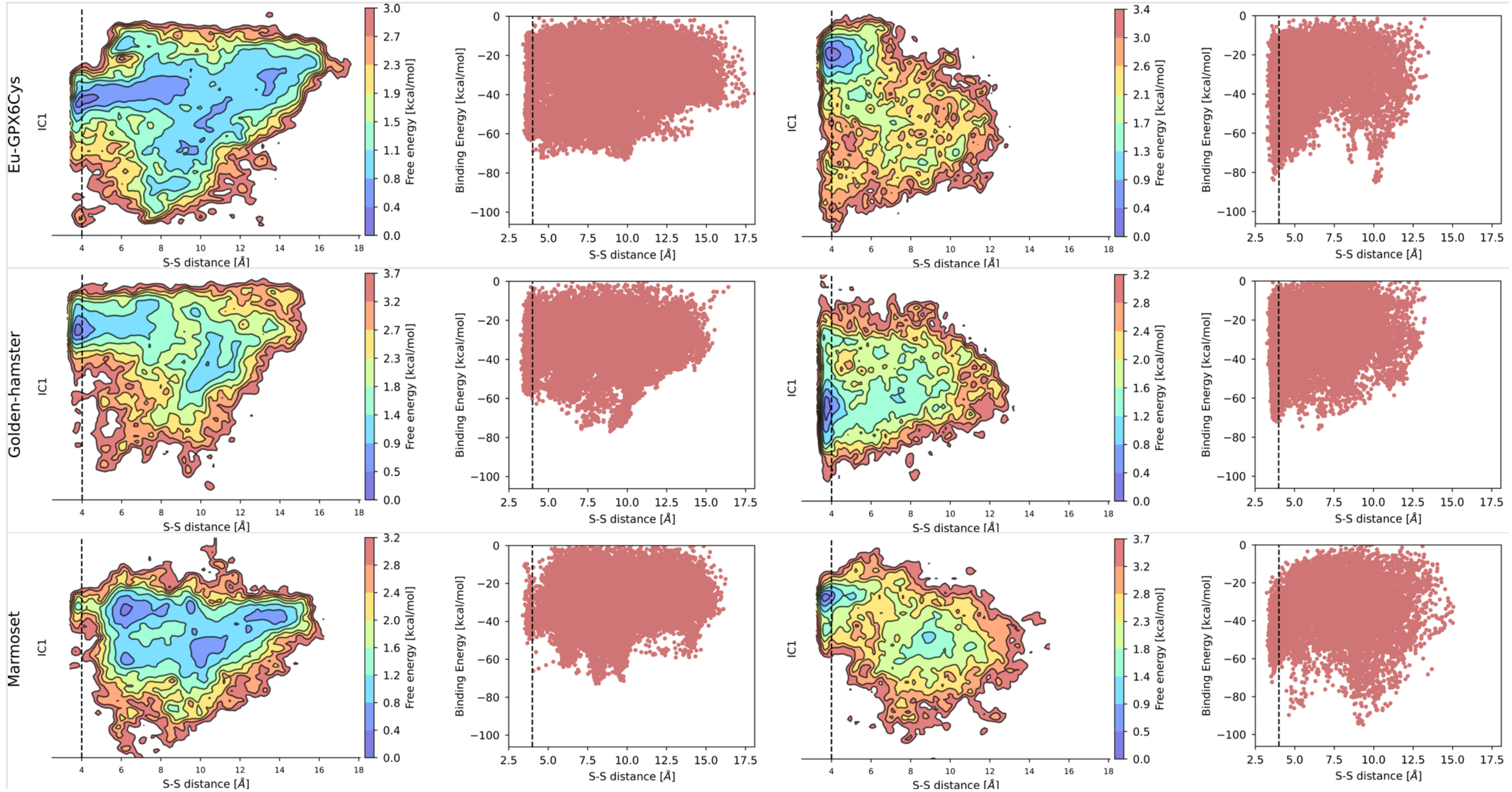

Eu-GPX6Cys
Golden-hamster
Marmoset
IC1
S-S distance [Å]
Free energy [kcal/mol]
Binding Energy [kcal/mol]

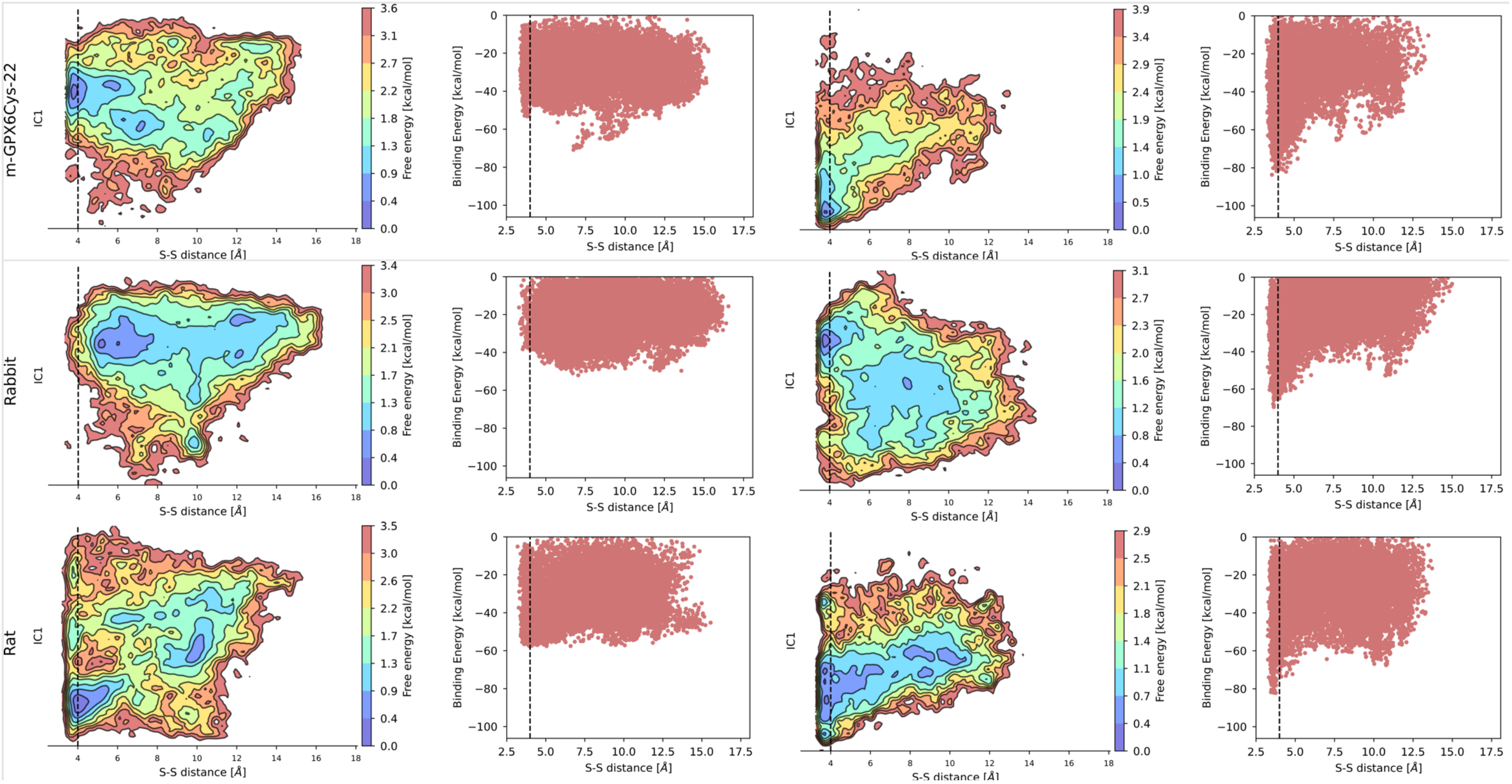

m-GPX6Cys-22
Rabbit
Rat
IC1
S-S distance [Å]
Free energy [kcal/mol]
Binding Energy [kcal/mol]

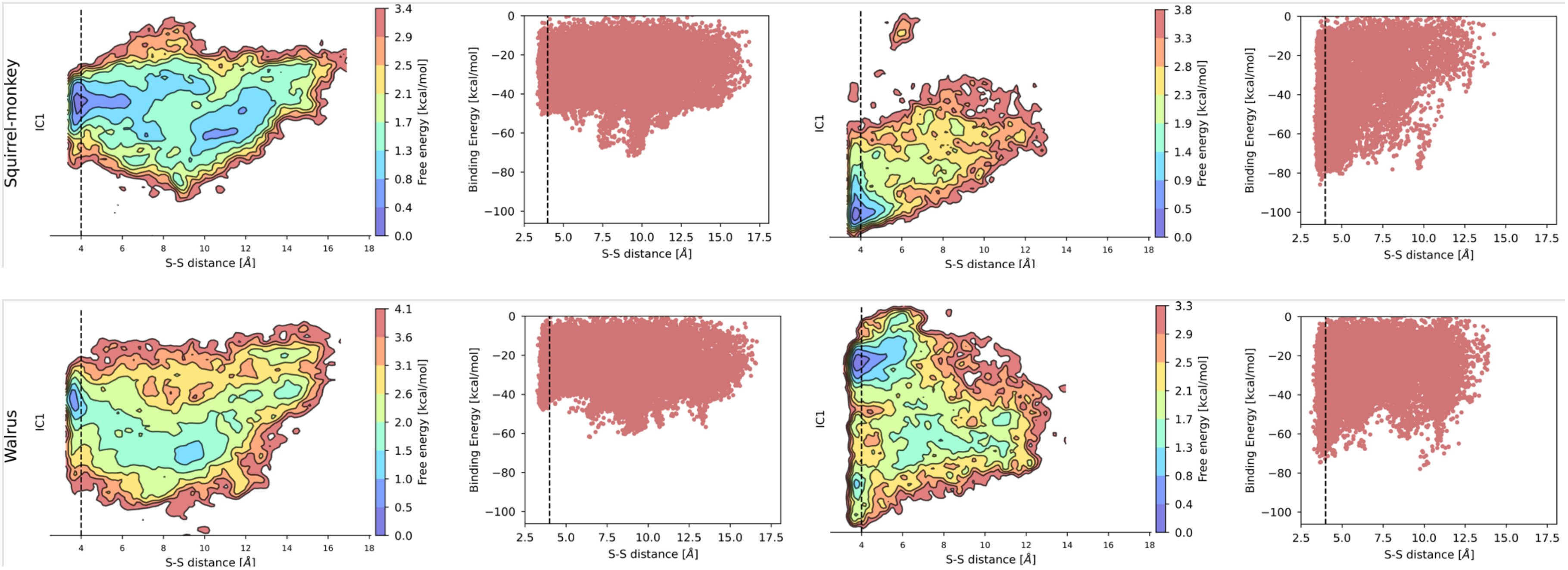


**Figure S14**. Free energy profiles for the docking of glutathione (left) and glutathione disulfide (right) to ancestral and modern GPX6$_{Cys}$ proteins. The x-axis represents the distance between the catalytic cysteine sulphur atom and the closest ligand's sulphur atom, while the Y-axis shows the slowest TICA coordinate or the binding free energy. The vertical dashed line represents a distance of 4Å.

### Tables

**Table S1**. dN/dS ratios for the GPX family and protein regions in lineages where GPX6 has Sec (Fig. 1, solid red branches), exchanged Sec for Cys (Fig. 1, dashed green branches) or inherited Cys (Fig. 1, solid green branches). In some lineages for which PAML estimates very few synonymous changes compared to the non-synonymous changes, unnaturally large dN/dS values can occur; these are marked with a #. The dN/dS ratio for all branches is the null hypothesis (one ratio for all branches) used in the likelihood ratio test contrasting the two previous ones. P-values are obtained based on a $\chi^2$ distribution with d.f=2. In bold, significant P-values.

| Protein | Region | dN/dS in branches where GPX6 has | | | | P-value |
|---|---|---|---|---|---|---|
| | | Sec | Cys after Sec was lost | Inherited Cys | All | |
| **GPX1$_{Sec}$** | Full length | 0.080 | 0.045 | 0.087 | 0.074 | 0.115 |
| | N-terminus | **0.043** | **0.009** | **0.190** | **0.034** | **0.046** |
| | GPX | 0.064 | 0.040 | 0.069 | 0.060 | 0.534 |
| | C-terminus | 0.085 | 0.052 | 0.114 | 0.081 | 0.328 |
| **GPX2$_{Sec}$** | Full length | **0.069** | **0.029** | **0.041** | **0.055** | **0.024** |
| | N-terminus | 0.032 | 0.001 | 0.001 | 0.032 | 0.999 |
| | GPX | 0.075 | 0.042 | 0.038 | 0.060 | 0.191 |
| | C-terminus | 0.055 | 0.017 | 0.048 | 0.043 | 0.100 |
| **GPX3$_{Sec}$** | Full length | 0.132 | 0.131 | 0.077 | 0.125 | 0.222 |
| | **N-terminus** | **0.241** | **0.038** | **0.461** | **0.181** | **0.022** |
| | GPX | 0.094 | 0.108 | 0.056 | 0.091 | 0.439 |
| | C-terminus | 0.105 | 0.195 | 0.058 | 0.114 | 0.161 |
| **GPX4$_{Sec}$** | Full length | 0.071 | 0.073 | 0.112 | 0.076 | 0.380 |
| | N-terminus | 0.108 | 0.018 | 0.123 | 0.082 | 0.264 |
| | GPX | **0.062** | **0.007** | **0.203** | **0.061** | **$1x10^{-4}$** |
| | C-terminus | 0.043 | 0.003 | 0.033 | 0.030 | 0.126 |
| **GPX5$_{Cys}$** | Full length | 0.294 | 0.258 | 0.429 | 0.305 | 0.061 |
| | N-terminus | 0.678 | 0.634 | 0.959 | 0.716 | 0.716 |
| | GPX | 0.233 | 0.145 | 0.219 | 0.212 | 0.227 |
| | C-terminus | 0.237 | 0.225 | 0.358 | 0.250 | 0.379 |

| | | | | | | |
|---|---|---|---|---|---|---|
| **GPX7**$_{Cys}$ | Full length | 0.141 | 0.086 | 0.157 | 0.137 | 0.377 |
| | N-terminus | 0.190 | #999 | 0.005 | 0.122 | 0.070 |
| | GPX | 0.083 | 0.080 | 0.117 | 0.088 | 0.712 |
| | C-terminus | 0.185 | 0.080 | 0.224 | 0.178 | 0.242 |
| **GPX8**$_{Cys}$ | Full length | 0.228 | 0.156 | 0.156 | 0.203 | 0.199 |
| | N-terminus | 0.169 | 0.195 | 0.112 | 0.158 | 0.775 |
| | GPX | 0.223 | 0.155 | 0.198 | 0.207 | 0.616 |
| | C-terminus | 0.217 | 0.161 | 0.104 | 0.194 | 0.486 |

**Table S2:** Foreground branches tested using the branch-site model, with the amino acid sites inferred as under selection using the Bayes Empirical Bayes inference listed when supported by a P-value < 0.05. P-values are obstained based on a $\chi^2$ distribution with d.f=1. Posterior probabilities of selection are shown in parehtneses, in bold face when P > 0.9. All foreground branches used here are lineages where Sec was exchanged for Cys (Fig 1, dashed green branches)

| Region | Foreground branches | P-value | Sites under selection |
|---|---|---|---|
| **GPX domain** | Eumuroida, Rabbit, Primate, Cat, Walrus | **0.046** | 45 (0.783); 46 (0.571); **50 (0.946); 52 (0.936); 56 (0.950);** 57 (0.513); **62 (0.995);** 63 (0.535); 70 (0.552); **74 (0.992);** 75 (0.573); **77 (0.975); 83 (0.995);** 91 (0.728); **110 (0.937); 126 (0.947)**; 143 (0.799); 149 (0.606) |
| **Full protein** | Eumuroida, Rabbit | **0.006** | 16 (0.741); **45 (0.913);** 56 (0.895); **74 (0.992)**; **77 (0.942)**; **83 (0.992)**; 126 (0.822); 171 (0.784); 214 (0.573); 215 (0.668) |
| | Eumuroida, Rabbit, Primate | **0.008** | **16 (0.988)**; 45 (0.858); **56 (0.935)**; 63 (0.501); 73 (0.503); **74 (0.991)**; **77 (0.971)**; **83 (0.988)**; **110 (0.925)**; **126 (0.937)**; 149 (0.554); 171 (0.757); 189 (0.520); 207 (0.502); 212 (0.758); 214 (0.698); 215 (0.682); 216 (0.822) |
| | Eumuroida, Rabbit, Primate, Cat, | 0.054 | |
| | Eumuroida, Rabbit, Primate, Walurs | 0.050 | |
| | Eumuroida, Rabbit, Primate, Cat, Walrus | 0.127 | |

**Table S3:** Convergent (sites change to the same amino acid) and pseudo-convergent (sites change to different amino acids) sites identified between GPX6$_{Cys}$ lineages by CONVERG2(Zhang and Kumar, *MBE* 1997). The two left-most columns are the branches between which convergent or pseudo-convergent sites are identified. The number gives the amino acid site where convergence is identified; the brackets represent the (ancestral amino acids, derived amino acids) in the order of the branches. Here, (SM-M) stands for the Squirrel monkey – marmoset internal branch, (GH-CH) stands for the Golden hamster – Chinese hamster internal branch and (rat-mouse) represents the rat-mouse internal branch. The branch names given in green indicate the branches upon which we have inferred the Sec-to-Cys exchange to have occurred. Convergent sites are in bold, all other sites are pseudo-convergent.

| | | | | | | | | | | | |
|---|---|---|---|---|---|---|---|---|---|---|---|
| (SM-M) | Eumuroida | 15 (GG,SA) | 166 (SK,NE) | | | | | | | | |
| (SM-M) | Rabbit | 15 (GG,SS) | 34 (GG, NE) | 62 (HH,PY) | 91 (FF, LL) | 110 (TK,AR) | 166 (SK,NR) | 212 (EE,AK) | | | |
| (SM-M) | Cat | 26 (NK,DT) | **48 (NN,SS)** | 62 (HH,PV) | 90 (NP,SR) | 91 (FF,LY) | 166 (SS,ND) | 190 (DD,HN) | | | |
| (SM-M) | Walrus | 91 (FF,LS) | | | | | | | | | |
| (SM-M) | Golden hamster | 48 (NN,ST) | | | | | | | | | |
| (SM-M) | Chinese hamster | 62 (HH,PY) | | | | | | | | | |
| (SM-M) | (Rat-mouse) | 48 (NN,SD) | 90 (QN,PS) | | | | | | | | |
| Eumuroida | Rabbit | 15 (GG,AS) | 45 (LL,KR) | **56 (KK,QQ)** | 83 (AA,TR) | 92 (GG,NS) | **143 (KK,NN)** | 165 (SS,TP) | 166(KK,ER) | **167 (QQ,HH)** | **171 (EE,DD)** |
| Eumuroida | Cat | **70 (TT,SS)** | **143 (KK,NN)** | 165 (SS,TP) | 166 (KS,ED) | 207(KK,QR) | | | | | |
| Eumuroida | Walrus | 165 (SS,TA) | | | | | | | | | |
| Eumuroida | Squirrel monkey | 48 (LL,KF) | **63 (VV,II)** | | | | | | | | |
| Eumuroida | Marmoset | **70 (TT,SS)** | | | | | | | | | |
| Eumuroida | Rat | 127 (FY,YF) | | | | | | | | | |
| Eumuroida | Chinese hamster | 27 (MA,AE) | | | | | | | | | |
| Eumuroida | (Rat-mouse) | 27 (MA,AS) | 45 (LK,KN) | 165 (ST,PT) | | | | | | | |
| Rabbit | Cat | 62 (HH,YV) | 91 (FF,LY) | **143 (KK,NN)** | **165 (SS,PP)** | 166 (KS,RD) | | | | | |
| Rabbit | Walrus | 51 (YY,DH) | 91 (FF,LS) | 165 (SP,PA) | | | | | | | |
| Rabbit | Squirrel monkey | 45 (LL,FR) | | | | | | | | | |
| Rabbit | Marmoset | 73 (GG,AS) | **208 (SS,AA)** | | | | | | | | |

| | | | | | |
|---|---|---|---|---|---|
| Rabbit | Mouse | 51 (YY,DF) | 200 (KQ,AH) | | |
| Rabbit | Rat | 36 (TT,SA) | 192 (VV,IA) | | |
| Rabbit | Golden hamster | 36 (TT,SA) | | | |
| Rabbit | Chinese hamster | **62 (HH,YY)** | 189 (PP,TS) | | |
| Rabbit | (Rat-mouse) | 11 (PP,LS) | 45 (LK,RN) | 160 (ST,PP) | 200 (KK,AQ) |
| Rabbit | (GH-CH) | 167 (QH,HY) | 171 (ED,DN) | | |
| Cat | Walrus | 91 (FF,YS) | 165 (SS,PA) | | |
| Cat | Squirrel monkey | 34 (KK,TN) | 50 (EE,GD) | | |
| Cat | Marmoset | **70 (TT,SS)** | | | |
| Cat | Golden hamster | 48 (NN,TS) | | | |
| Cat | Chinese hamster | 62 (HH,YV) | | | |
| Cat | (Rat-mouse) | 48 (NN,SD) | 90 (PQ,RP) | 165 (ST,PP) | |
| Walrus | Mouse | 51 (YY,HF) | 54 (QQ,PN) | | |
| Walrus | (Rat-mouse) | 165 (ST,AP) | | | |
| Walrus | (GH-CH) | **54 (QQ,PP)** | | | |
| Squirrel monkey | (Rat-mouse) | 45 (LK,FN) | | | |
| Marmoset | Rat | 94 (IT,TS) | | | |
| Marmoset | (Rat-mouse) | **94 (II,TT)** | | | |
| Mouse | (Rat-mouse) | 200 (QK,HQ) | | | |
| Mouse | (GH-CH) | 54 (QQ,NP) | | | |
| Rat | Golden hamster | **36 (TT,AA)** | | | |
| Rat | (Rat-mouse) | 94 (TI,ST) | 205 (IT,VI) | | |
| Golden hamster | (Rat-mouse) | 48 (NN,TD) | | | |
| Chinese hamster | (Rat-mouse) | 27 (AA,ES) | | | |

**Table S4:** Convergent and pseudoconvergent sites in GPX1 between the GPX$6_{Cys}$ lineages, where the sequences were also available, as identified by CONVERG2(Zhang and Kumar, *MBE* 1997). The two left-most columns are the branches between which convergent or pseudo-convergent lineages are identified. The number gives the amino acid site where convergence is identified; the brackets represent the (ancestral amino acids, derived amino acids) in the order of the branches. Here, (SM-M) stands for the Squirrel monkey – marmoset internal branch, (GH-CH) stands for the Golden hamster – Chinese hamster internal branch and (rat-mouse) represents the rat-mouse internal branch. The branch names given in green indicate the branches upon which we have inferred the Sec-to-Cys exchange in GPX$6_{Cys}$ to have occurred. Strict convergent sites are given in bold, all other sites are pseudo-convergent.

| | | | |
|---|---|---|---|
| Rabbit | Mouse | **177 (PP,SS)** | |
| Rabbit | Rat | **138 (AA,SS)** | **175 (QQ,KK)** |
| Rabbit | Golden hamster | **10 (SS,NN)** | 41 (RK,ER) |
| Cat | (Mouse-Rat) | 108 (EE,QN) | |
| Rat | Golden hamster | **4 (TT,AA)** | |

**Table S5:** Convergent and pseudoconvergent sites in GPX2 between the GPX$6_{Cys}$ lineages, where the sequences were also available, as identified by CONVERG2(Zhang and Kumar, *MBE* 1997). The two left-most columns are the branches between which convergent or pseudo-convergent lineages are identified. The number gives the amino acid site where convergence is identified; the brackets represent the (ancestral amino acids, derived amino acids) in the order of the branches. Here, (SM-M) stands for the Squirrel monkey – marmoset internal branch, (GH-CH) stands for the Golden hamster – Chinese hamster internal branch and (rat-mouse) represents the rat-mouse internal branch. The branch names given in green indicate the branches upon which we have inferred the Sec-to-Cys exchange in GPX$6_{Cys}$ to have occurred. Strict convergent sites are given in bold, all other sites are pseudo-convergent.

| | | |
|---|---|---|
| Rabbit | Mouse | 47 (EQ,QE) |

**Table S6:** Convergent and pseudoconvergent sites in GPX3 between the GPX$6_{Cys}$ lineages, where the sequences were also available, as identified by CONVERG2(Zhang and Kumar, *MBE* 1997). The two left-most columns are the branches between which convergent or pseudo-convergent lineages are identified. The number gives the amino acid site where convergence is identified; the brackets represent the (ancestral amino acids, derived amino acids) in the order of the branches. Here, (SM-M) stands for the Squirrel monkey – marmoset internal branch, (GH-CH) stands for the Golden hamster – Chinese hamster internal branch and (rat-mouse) represents the rat-mouse internal branch. The branch names given in green indicate the branches upon which we have inferred the Sec-to-Cys exchange in GPX$6_{Cys}$ to have occurred. Here, the marmoset branch is given as a branch where the Sec-to-Cys exchange in GPX$6_{Cys}$ has been estimated, given that the squirrel monkey sequence is unavailable for this protein. Strict convergent sites are given in bold, all other sites are pseudo-convergent.

| | | | |
|---|---|---|---|
| Marmoset | Eumuroida | **5 (VV,MM)** | 172 (SA,LS) |
| Marmoset | Rabbit | 33 (VI,LV) | |
| Marmoset | Cat | 5 (VV,MG) | |
| Marmoset | (GH-CH) | 33 (VI,LV) | |
| Eumuroida | Rabbit | 135 (VV,IM) | |
| Eumuroida | Cat | 5 (VV,MG) | |
| Rabbit | Cat | **126 (GG,NN)** | **152 (II,VV)** |
| Rabbit | Chinese hamster | 152 (II,VK) | |
| Rabbit | (GH-CH) | **33 (II,VV)** | **107 (FF,VV)** |
| Cat | Mouse | 154 (II,LV) | |
| Cat | Chinese hamster | 152 (II,VK) | |
| Cat | (GH-CH) | 154 (II,LV) | |
| Mouse | (GH-CH) | **154 (II,VV)** | |

**Table S7.** Convergent and pseudoconvergent sites in GPX4 between the $GPX6_{Cys}$ lineages, where the sequences were also available, as identified by CONVERG2(Zhang and Kumar, *MBE* 1997).

No convergent sites found.

**Table S8:** Convergent and pseudoconvergent sites in GPX5 between the $GPX6_{Cys}$ lineages, where the sequences were also available, as identified by CONVERG2(Zhang and Kumar, *MBE* 1997). The two left-most columns are the branches between which convergent or pseudo-convergent lineages are identified. The number gives the amino acid site where convergence is identified; the brackets represent the (ancestral amino acids, derived amino acids) in the order of the branches. Here, (SM-M) stands for the Squirrel monkey – marmoset internal branch, (GH-CH) stands for the Golden hamster – Chinese hamster internal branch and (rat-mouse) represents the rat-mouse internal branch. The branch names given in green indicate the branches upon which we have inferred the Sec-to-Cys exchange in $GPX6_{Cys}$ to have occurred. Strict convergent sites are given in bold, all other sites are pseudo-convergent.

| | | | |
|---|---|---|---|
| (SM-M) | Rabbit | 140 (RR,QL) | |
| (SM-M) | Cat | 5 (KK,RN) | **140 (RR,QQ)** |
| (SM-M) | Walrus | 18 (AT,MS) | **29 (QQ,RR)** |
| (SM-M) | Squirrel monkey | 151 (LV,VE) | |
| (SM-M) | Marmoset | 145 (SL,LI) | |
| (SM-M) | Mouse | 151 (LL,VM) | |
| (SM-M) | Golden hamster | 5 (KK,RT) | |
| (SM-M) | Chinese hamster | 151 (LL,VM) | |
| (SM-M) | (GH-CH) | 18 (AS,MA) | 148 (TS,AA) |
| Eumuroida | Rabbit | 48 (AA,IS) | |
| Eumuroida | Cat | **130 (DD,NN)** | |
| Eumuroida | Mouse | 130 (DN,ND) | |
| Eumuroida | Rat | 52 (SL,LT) | |

| | | | | |
|---|---|---|---|---|
| Eumuroida | Golden hamster | 17 (FL,LF) | 48 (AI,IM) | 103 (AV,VA) |
| Eumuroida | (Mouse-rat) | 109 (SY,YF) | | |
| Rabbit | Cat | 82 (EE,GK) | 140 (RR,LQ) | |
| Rabbit | Squirrel monkey | 0 (KQ,QK) | | |
| Rabbit | Rat | **13 (DD,NN)** | 86 (KK,NE) | |
| Rabbit | Golden hamster | 0 (KK,QR) | 48 (AI,SM) | |
| Rabbit | (Mouse-rat) | **65 (GG,KK)** | **67 (YY,FF)** | |
| Cat | Mouse | 83 (KK,TN) | 130 (DN,ND) | |
| Cat | Golden hamster | 5 (KK,NT) | | |
| Walrus | Marmoset | 21 (GK,EE) | | |
| Walrus | (Mouse-rat) | **26 (QQ,PP)** | | |
| Walrus | (GH-CH) | 18 (TS,SA) | | |
| Squirrel monkey | Mouse | 151 (VL,EM) | | |
| Squirrel monkey | Golden hamster | 0 (QK,KR) | | |
| Squirrel monkey | Chinese hamster | 151 (VL,EM) | | |
| Marmoset | Chinese hamster | **94 (SS,AA)** | | |
| Mouse | Chinese hamster | **151 (LL,MM)** | | |
| Mouse | (Mouse-rat) | 104 (TS,MT) | 155 (NK,SN) | |
| Golden hamster | (GH-CH) | 101 (IT,VI) | | |

**Table S9:** Convergent and pseudoconvergent sites in GPX7 between the $GPX6_{Cys}$ lineages, as identified by CONVERG2(Zhang and Kumar, *MBE* 1997).. The two left-most columns are the branches between which convergent or pseudo-convergent lineages are identified. The number gives the amino acid site where convergence is identified; the brackets represent the (ancestral amino acids, derived amino acids) in the order of the branches. Here, (SM-M) stands for the Squirrel monkey – marmoset internal branch, (GH-CH) stands for the Golden hamster – Chinese hamster internal branch and (rat-mouse) represents the rat-mouse internal branch. The branch names given in blue indicate the branches upon which we have inferred the Sec-to-Cys exchange in $GPX6_{Cys}$ to have occurred. Here, the marmoset branch is given as a branch where the Sec-to-Cys exchange in $GPX6_{Cys}$ has been estimated, given that the squirrel monkey sequence is unavailable for this protein Strict convergent sites are given in bold, all other sites are pseudo-convergent.

| | | |
|---|---|---|
| Marmoset | Mouse | 95 (AA,SD) |
| Marmoset | (GH-CH) | 95 (AA,SD) |
| Eumuroida | Rabbit | **109 (SS,PP)** |
| Eumuroida | Chinese hamster | 116 (HR,RQ) |
| Rabbit | Mouse | 30 (HY,RH) |
| Mouse | (GH-CH) | **49 (SS,TT)** **95 (AA,DD)** 111 (EE,AQ) |

**Table S10:** Convergent and pseudoconvergent sites in GPX8 between the $GPX6_{Cys}$ lineages, as identified by CONVERG2(Zhang and Kumar, *MBE* 1997).. The two left-most columns are the branches between which convergent or pseudo-convergent lineages are identified. The number gives the amino acid site where convergence is identified; the brackets represent the (ancestral amino acids, derived amino acids) in the order of the branches. Here, (SM-M) stands for the Squirrel monkey – marmoset internal branch, (GH-CH) stands for the Golden hamster – Chinese hamster internal branch and (rat-mouse) represents the rat-mouse internal branch. The branch names given in blue indicate the branches upon which we have inferred the Sec-to-Cys exchange in $GPX6_{Cys}$ to have occurred. Strict convergent sites are given in bold, all other sites are pseudo-convergent.

| | | |
|---|---|---|
| Eumuroida | (GH-CH) | 12 (LF,FY) |
| Rabbit | (GH-CH) | 121 (VI,IV) |
| Mouse | Chinese hamster | 59 (KK,QR) |
| Rat | (Mouse-rat) | 18 (QL,EQ) |
| Chinese hamster | (GH-CH) | 51 (MK,TM) |
